# A novel deep learning-based framework reveals a continuum of chromatin sensitivities across transcription factors

**DOI:** 10.1101/2025.08.11.669605

**Authors:** Lukas Burger, Noa Gil, Dirk Schübeler

## Abstract

The genome-wide binding of many transcription factors (TFs) depends not only on the presence of their recognition motifs, but also on the surrounding chromatin context. This raises the question of how binding specificity is achieved, which is essential for the proper activation of regulatory regions during developmental and differentiation processes. Two competing hypotheses have been proposed: either opening is a hierarchical process, whereby chromatin-insensitive TFs bind initially and make the DNA accessible for other TFs; or multiple chromatin-sensitive TFs bind cooperatively, enabling them to overcome the chromatin barrier. Here, using complementary genomic- and deep-learning-based techniques, we provide a quantitative framework to assess chromatin sensitivity of individual TFs, and hone in on the definitions and implications of such sensitivity. Our findings demonstrate that the degree of chromatin sensitivity is a continuum across TFs, showing that regulatory regions are activated neither by a purely hierarchical nor purely cooperative mechanism, but by a mixture of both mechanisms. The presented framework allows for a locus-specific as well as global, quantitative assessment of chromatin sensitivity and should help in understanding the binding behaviour of individual TFs as well as their potential contribution to gene regulation.

---

Gene expression is regulated by transcription factors (TFs), which are DNA-binding proteins that recognize short recognition sequences (‘motifs’) in the DNA and, upon binding, activate or repress transcription. Regulatory regions that drive gene expression, such as promoters and enhancers, often contain binding sites for multiple TFs (Wang et al., 2012; Yan et al., 2013). However, for most TFs, the binding sites in these regions comprise a small subset of all potential binding sites and only a fraction of all motifs in the genome are generally bound. This indicates that the presence of a motif alone is often not sufficient to induce binding, but that the surrounding genomic context also plays a role in shaping binding patterns (Wang et al., 2012). Genomic context that is refractory to binding may include the presence of nucleosomes, of nucleosomes with particular modifications, or of epigenetic modifications of the DNA itself, such as DNA methylation (Arvey et al., 2012; Cirillo et al., 2002; Yin et al., 2017).

To describe the degree to which a TF’s binding is dependent on the genomic context, we have recently introduced the term ‘chromatin sensitivity’ (Isbel et al., 2022). The finding that most TFs bind only a small fraction of their motifs suggests strong chromatin sensitivity for most TFs. However, this raises the question of how previously-unbound regulatory regions become bound during development and cellular differentiation. At least two potential mechanisms have been proposed. The first suggests that the opening of regulatory regions is a hierarchical process: certain TFs act as so-called pioneer factors, which are capable of binding their motifs even in closed chromatin, initiating the opening of chromatin around their binding sites and subsequently enabling the binding of other TFs to the now permissive chromatin. This model is often invoked to explain the process of lineage differentiation, which is commonly initiated by the expression of one or a few specific TFs (e.g. Boller et al., 2016; Cirillo et al., 2002; Mayran et al., 2018; Minderjahn et al., 2020). Alternatively, it has been suggested that TFs can cooperate with one another to be able to bind their respective sites in closed regulatory regions, where each TF individually would be unable to bind (Kim & Wysocka, 2023). Such cooperativity can be mediated via protein-protein interactions, or can be nucleosome or DNA-induced (Jolma et al., 2015; Mirny, 2010; Thanos & Maniatis, 1995).

The ability of each model to explain TF binding and regulatory activity is subject to debate, fuelled in part by flexibly defined concepts and seemingly contradictory findings (Bulyk et al., 2023; Stoeber et al., 2024). For example, some TFs are being referred to as pioneers because of a demonstrated ability to bind in closed chromatin, but they have also been shown to bind only a small fraction of their total predicted motifs, or to exhibit cell-type specific binding behaviour, both indicating that their binding is not unrestricted by chromatin (Donaghey et al., 2018; Michael et al., 2020; Minderjahn et al., 2020; Pham et al., 2013). Relatedly, the binding patterns of some TFs have been shown to be strongly influenced by TF-extrinsic properties, such as differential affinity to motif variants or a TF’s expression level (Blassberg et al., 2022; Gibson et al., 2024; Minderjahn et al., 2020; Pham et al., 2013). Combined, such findings may highlight the need for more precise definitions of both pioneering and chromatin sensitivity.

Despite these ambiguities, discerning the cooperative and hierarchical models could substantially promote our understanding of TF binding determinants and thus of gene expression regulation. One approach to tackling this is to employ the fact that each model makes strong predictions regarding the distribution of chromatin sensitives across TFs: While the cooperativity model suggests that most TFs are chromatin-sensitive (possibly to varying degrees), the pioneering model predicts that there are two classes of TFs, a large group of chromatin-sensitive TFs and a small group of chromatin-insensitive pioneers which can bind their motifs in the genome irrespective of context. Accordingly, developing a quantitative framework to assess chromatin sensitivities across TFs should allow an evaluation of the validity of each model, as well as indicate the degree to which they need to be modified to incorporate TF-extrinsic properties.

Using genomic approaches, we here show that chromatin sensitivity is not binary, but rather continuous and generally dependent on motif strength. Based on these insights, we provide a quantitative framework of chromatin sensitivity, which builds on the deep learning tool Enformer (Avsec et al., 2021), and assesses chromatin sensitivity as the degree to which binding is determined by the binding motif sequence versus the flanking sequence around the motif, the latter indicating to what degree genomic context modulates binding. Application of this approach to many mouse and human datasets reveals that chromatin sensitivity is a continuum across TFs. While this disagrees with the binary classification of the pioneering model, it also argues against a purely cooperative model. Our results therefore suggest a mixed model of both cooperative and pioneer-like binding behaviour.

## Results

### Chromatin sensitivity is not binary and generally depends on motif strength

According to a definition we previously proposed, chromatin sensitivity refers to the sensitivity of a TF’s binding patterns to the presence of nucleosomes, nucleosomes marked with particular chromatin modifications, or to epigenetic modifications of the DNA, such as DNA methylation (Isbel et al., 2022). An assessment of chromatin sensitivity therefore requires a comparison of where a TF binds in the genome to where it would bind if the DNA was freely accessible and devoid of any such chromatin characteristics. Based on the hypothesis that the latter can be approximated as all the genomic loci with a predicted binding motif, we previously hypothesized that chromatin sensitivity can be assessed as the fraction of bound strong motifs in the genome: chromatin-insensitive TFs should be able to bind a large majority of these motifs, whereas chromatin-sensitive TFs only a small fraction (Figure 1a, Isbel et al. (2022)). To investigate this idea, we started out by making use of and expanding a collection of ChIP-seq datasets from mouse embryonic stem cells (ESCs) that we had previously manually curated (Ginno et al., 2020). These datasets were uniformly processed and carefully selected to have good enrichments and little to no GC and open chromatin biases, which can strongly influence binding profiles (Benjamini & Speed, 2012; Teng & Irizarry, 2016; Teytelman et al., 2013). In its final version, this collection of datasets includes ChIP-seq data of 27 TFs (Table 1). For each of these TFs, we predicted binding motifs genome-wide using their weight matrices as inferred from the 500 strongest peaks (Ginno et al. (2020), Supplementary Figure 1). The strength of each predicted motif was quantified with the commonly used log-odds score, which is an estimate of relative affinity (Berg & von Hippel, 1988).

**Figure 1.**
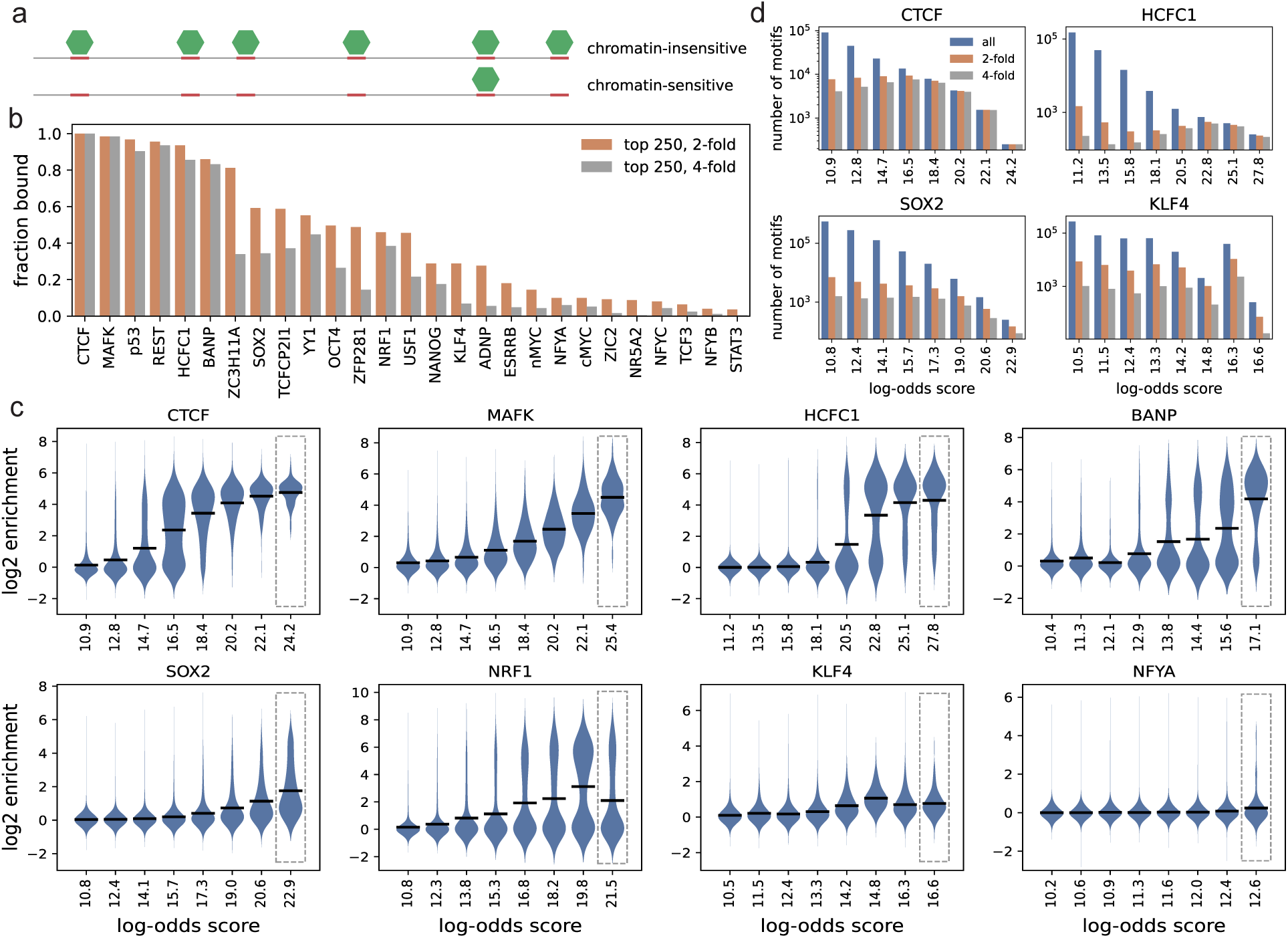
(a) TFs (green hexagons) that bind all of their strong motifs (red rectangles) are chromatin-insensitive whereas those that do not are chromatin-sensitive. (b) Fraction of strong motifs that are bound for each TF. Number of motifs selected and enrichment cut-offs indicated in the legend. TFs were sorted by decreasing fraction of at least 2-fold bound top 250 motifs. (c) Binding enrichment in bins of increasing log-odds score for a selected set of TFs, sorted as in b. The top bin corresponds to the top 250 motifs as in b (grey dashed rectangle), while all other motifs were binned into equal-width log-odds score bins. Bin midpoints are indicated. Black lines indicate mean enrichments per bin. To facilitate visual comparisons, violins for each bin were scaled to have the same width. (d) Total number of motifs as well as total number of 2-fold and 4-fold bound motifs as a function of log-odds score for four example TFs. Log-odds score bins as in c.

**Table 1.** Data-sets for mouse ESCs.

| TF | GEO accession (IP, control) | reference |
| --- | --- | --- |
| p53 | GSM647224, GSM647222 | (M. Li et al., 2012) |
| ZIC2 | GSM1499117, GSM1499114 | (Luo et al., 2015) |
| ZFP281 | GSM2141217, GSM2141216 | (Fidalgo et al., 2016) |
| NR5A2 | GSM2429232, GSM1003811 | (Atlasi et al., 2019; Yue et al., 2014) |
| YY1 | GSM3615268, GSM1003811 | (Atlasi et al., 2019; Yue et al., 2014) |
| USF1 | GSM3483911, GSM3483832 | (Scelfo et al., 2019) |
| TCFCP2I1 | GSM288350, GSM288358 | (Chen et al., 2008) |
| STAT3 | GSM288353, GSM288358 | (Chen et al., 2008) |
| nMYC | GSM288357, GSM288358 | (Chen et al., 2008) |
| TCF3 | GSM307143, GSM307154 | (Marson et al., 2008) |
| REST | GSM671095, GSM671103 | (Arnold et al., 2013) |
| NRF1 | GSM1891642, GSM1891643 | (Domcke et al., 2015) |
| NFYA | GSM1370111, GSM1370110 | (Oldfield et al., 2014) |
| NFYB | GSM1370112, GSM1370110 | (Oldfield et al., 2014) |
| NFYC | GSM1370113, GSM1370110 | (Oldfield et al., 2014) |
| HCFC1 | GSM1003799, GSM1003811 | (Yue et al., 2014) |
| MAFK | GSM1003809, GSM1003811 | (Yue et al., 2014) |
| ZC3H11A | GSM1003810, GSM1003811 | (Yue et al., 2014) |
| OCT4 | GSM2417142, GSM2417127 | (Chronis et al., 2017) |
| SOX2 | GSM2417143, GSM2417127 | (Chronis et al., 2017) |
| KLF4 | GSM2417144, GSM2417127 | (Chronis et al., 2017) |
| cMYC | GSM2417145, GSM2417127 | (Chronis et al., 2017) |
| NANOG | GSM2417187, GSM2417127 | (Chronis et al., 2017) |
| ESRRB | GSM2417188, GSM2417127 | (Chronis et al., 2017) |
| CTCF | GSM747535, GSM747546 | (Stadler et al., 2011) |
| BANP | GSM4708466, GSM4708472 | (Grand et al., 2021) |
| ADNP | GSM2582357, GSM2582354 | (Ostapcuk et al., 2018) |

To determine the fraction of bound motifs for each TF, we selected the top 250 motifs with highest log-odds score, and defined bound as either a 2-fold or a 4-fold enrichment over control in a 200bp window around the motif (Figure 1b). Interestingly, this revealed a continuum of values across TFs. While some TFs, such as CTCF and MAFK, bind almost all of their top motifs, suggesting they are chromatin-insensitive according to the definition proposed above, other TFs, such as NFYA, bind below 10 percent of their top motifs, indicating they are likely chromatin-sensitive. Additionally, there is a substantial group of TFs that bind between 10-60 percent of their top motifs. These TFs, which include for example SOX2 and NRF1, can therefore not easily be assigned to either category of chromatin sensitivity. It is unclear how to interpret these intermediate fractions of bound sites - they may represent generally weak binding signal close to the cut-off, or reflect the existence of both a bound and an unbound sub-group. To distinguish between these two possibilities, we visualized the distribution of binding enrichments at these top 250 sites (right-most log-odds score bin (grey dashed rectangle) for each TF in Figure 1c and Supplementary Figure 2). This revealed either bimodal (e.g. NRF1) or heavy-tailed (e.g. SOX2) distributions of enrichments, indicating the presence of both bound and unbound sub-groups, and suggesting that TFs with an intermediate fraction of bound top sites are in fact chromatin-sensitive at their top motifs. Interestingly, despite binding more than 80 percent of their top sites (Figure 1b), HCFC1 and BANP also display bimodal distributions of enrichments at their top motifs, suggesting that they are chromatin-sensitive as well. Finally, TFs with a small fraction of bound top sites, such as NFYA, generally show a distribution of enrichments centered around 0, reflecting unbound sites, with a very small heavy tail towards high enrichments, representing bound sites (gray box in Figure 1c and Supplementary Figure 2), and are thus also chromatin-sensitive at these motifs.

The predicted chromatin sensitivity of SOX2, and equally so for OCT4 (Supplementary Figure 2), is surprising, as both TFs have been classified as pioneer factors based on their ability to bind in closed chromatin (Soufi et al., 2012, 2015). As the most strongly bound regions of SOX2 and OCT4 are generally co-bound by both proteins (Chen et al., 2008; Michael et al., 2020), in line with the inferred weight matrices containing the recognition sequences of both OCT4 and SOX2 (Supplementary Figure 1), this suggests that not even the joint binding of these two proteins is sufficient to overcome chromatin. In line with this finding, it was recently shown that OCT4 binding is strongly reduced when a nucleosome is positioned over the combined OCT4-SOX2 motif (Grand et al., 2024).

While informative, binding patterns at the top 250 motifs give only a very limited picture of genome-wide binding behaviours. In fact, the number of predicted motifs increases very rapidly with decreasing log-odds score, and the top 250 motifs generally represent only a minority of bound motifs (Figure 1d). Extending the above analysis to include predicted motifs with lower log-odds score reveals a much more complex picture of binding behaviour and thus of chromatin sensitivity (Figure 1c and Supplementary Figure 2). In general, for TFs with intermediate to high fractions of top bound motifs, average binding enrichments generally increase with increasing motif strength (black horizontal lines for first six TFs in Figure 1c, first 15 TFs in Supplementary Figure 2). The variability in binding within log-odd score bins, on the other hand, follows a more complicated pattern. For CTCF, while average binding (black horizontal lines) starts to rise above background around log-odds scores of 12.8, most of these motifs are unbound, likely because of their very low affinity. For motifs with larger scores up to around 16.5, both average binding and variability in binding increase with a steadily increasing fraction of bound motifs. For even larger log-odds scores, average binding still increases, but variability in binding starts to decrease again, so that in the highest log-odds score bins almost all motifs are bound. These observations suggest a motif-strength-dependent definition of chromatin sensitivity: while motif strength sets the average binding enrichment, the variability in binding at that strength (relative to the total variability in binding at all motifs) reflects the degree of chromatin sensitivity. In a given bin, motifs that are more strongly bound than the average likely lie in a permissive chromatin environment, whereas motifs that are more weakly bound in a more refractory environment. According to this logic, chromatin sensitivity for CTCF first increases, is largest at motifs around scores of 16.5, then decreases again and is likely small at the top motifs. MAFK, on the other hand, shows generally moderate variability in binding within fixed log-odds score bins, with very little change in variability for log-odds scores around 16.5 and higher, where binding enrichments start to rise above background, suggesting it is moderately chromatin-sensitive independent of motif strength. For HCFC1 and BANP, from log-odds scores 20.5 and 13.8 and on, where average binding levels start to rise above background, binding profiles look similar to CTCF at low and intermediate log-odds scores, with variability and thus chromatin sensitivity first increasing and then decreasing, but not down to levels comparable to CTCF at strong motifs, reflecting that both TFs are chromatin sensitivity also at their strongest motifs. For SOX2, average binding starts to rise at log-odds scores around 19 and continuously increases, with a concomitant increase in binding variability, indicating that SOX2 is most chromatin-sensitive at its strongest motifs. While NRF1 shows a similar behaviour, the fraction of bound sites appears to decrease from the second highest to the highest log-odds score bin, a behaviour we discuss in detail below. Finally, for KLF4 and NFYA, there is little to no relationship between binding and log-odds score, with average binding levels never clearly rising above background. As it is generally not clear whether degenerate version of low-complexity motifs (e.g. the motif of NFYA) still represent valid binding sites, it is difficult to interpret the binding behaviour of these TFs beyond their behaviour at their top motifs. Of note, it is generally unclear how accurate lower log-odds scores are, and in addition, in some cases very low correlations between motif strength and binding may also indicate that the inferred weight matrices used to call motifs (Supplementary Figure 1) are imprecise. These points will be discussed in detail below.

An implicit assumption when inferring chromatin sensitivity based on a comparison of binding variability across log-odds score bins is that chromatin contexts are roughly equally represented in the different bins. We wanted to check whether that is indeed the case, and as an indicator of chromatin context chose to focus on DNaseI signal (Hesselberth et al., 2009), which measures the degree to which the DNA is accessible, and has previously been shown to have largest predictive power among a variable set of chromatin features (Arvey et al., 2012). More specifically, we looked at DNaseI hypersensitive sites (DHSs), i.e. peaks of DNaseI signal. A potential complication in using DHSs directly, however, is that many TFs themselves create a DHS when binding to the DNA, making it difficult to distinguish accessibility created by the binding of the TF in question from the accessibility created by other TFs binding in the surrounding region.

While methods have been proposed to deconvolve the accessibility signal of multiple TFs (Pique-Regi et al., 2011; Sherwood et al., 2014), we opted for a simpler approach, by separating DHSs by their size: Comparing widths of DHSs at promoters, which are usually bound by multiple TFs, to strong distal binding sites of CTCF, where CTCF is thought to bind by itself (Fu et al., 2008), suggests that a cut-off of 300 nts roughly separates binding sites of a single TF and regions bound by multiple factors (Supplementary Figure 3). DHSs larger than 300 nts (’large DHSs’) thus likely represent a reasonable proxy for the presence of accessible chromatin around a TF’s motif independent of the TF in question.

Returning to the question of whether different chromatin contexts are equally represented across log-odds bins, we quantified the fraction of motifs in each log-odds score bin that overlap large DHSs (Supplementary Figure 4). This revealed that for most TFs, stronger motifs are more likely to overlap with large DHSs, suggesting among other things that motif variants are not randomly distributed in the genome (see below and Discussion).

For a subset of TFs (HCFC1, BANP, YY1, and NRF1), this effect is very strong, with the fraction of motifs overlapping large DHSs increasing by up to 50% from the weakest to the strongest motifs. The high overlap with large DHSs at the top sites of HCFC1 and BANP likely explains the large fraction of bound top motifs for these TFs (Figure 1b) despite their chromatin sensitivity at these sites (Figure 1c). In some cases, certain log-odds score bins exhibit a larger overlap with large DHS than expected by a generally increasing trend, which appears to have a direct effect on the amount of binding variability as well as average enrichments. For example, for NRF1, the second largest log-odds score bin exhibits larger variability in binding and higher average enrichments than the top log-odds bin (Figure 1c), likely due to the larger overlap with large DHSs for the former (Supplementary Figure 4).

The unequal distribution of chromatin states across log-odd score bins makes it difficult to assess how reliably binding variability (as well as average enrichments) can be compared across bins. We wondered whether chromatin sensitivity could be more accurately quantified by stratifying not only by motif strength but also directly by chromatin state, i.e. overlap with large DHSs. According to this logic, a TF would be considered to be chromatin-insensitive at motifs of a given strength if it binds at such motifs irrespective of a permissive chromatin environment, represented by the presence of large DHSs. Conversely, for a chromatin-sensitive TF the binding patterns would depend on the overlap with large DHSs (Figure 2a). To test this directly, we stratified the binding versus log-odds score curves (Figure 1c and Supplementary Figure 2) by the presence of large DHSs (Figure 2b and Supplementary Figure 5). This revealed that the presence of large DHSs is generally correlated with a shift to higher binding enrichments for all TFs, in agreement with large DHSs representing a permissive chromatin environment. For CTCF, the presence of large DHSs has a small effect on binding at the weakest and strongest motifs, while it strongly increases the fraction of motifs with high enrichments for motifs of intermediate strength, similar to the increased variability in binding at these motifs compared to weak and strong motifs in the previous analysis (Figure 1c). This likely indicates, as with the previous approach, chromatin insensitivity at the strongest motifs and chromatin sensitivity at intermediate motifs. Generally, the binding behaviour of CTCF stratified by large DHSs and log-odd scores suggests a new definition of chromatin sensitivity, closely related to the previous one: motif-strength-dependent chromatin sensitivity is reflected in the difference in average binding at motifs of a given strength within large DHS, i.e. within permissive chromatin, versus outside of large DHS, i.e. within non-permissive chromatin. Applying this concept across all TFs reveals a (at least qualitatively) similar picture as using the previous method: MAFK is moderately chromatin-sensitive independent of motif strength, for HCFC1 and BANP chromatin sensitivity first increases and then decreases, and for SOX2 and NRF1 chromatin sensitivity increases with increasing motif strength. Whereas the previous approach did not provide a clear picture for KLF4 and NFYA (except for possibly at their top motifs), stratification by large DHSs reveals increasing binding signal as a function of motif strength within large DHSs and largely unbound motifs outside of large DHSs, suggesting weakly increasing chromatin sensitivity as a function of motif strength. This behaviour can be seen for many TFs with low fractions of top-bound motifs (Supplementary Figure 5). In sum, while stratification by large DHSs likely results in a more quantitative assessment, as it circumvents the problem of unequal representation of large DHS across log-odds score bins, both genomic approaches reveal similar patterns of chromatin sensitivity and generally show that chromatin sensitivity is not binary, but rather a continuous measure that depends on motif strength.

**Figure 2.**
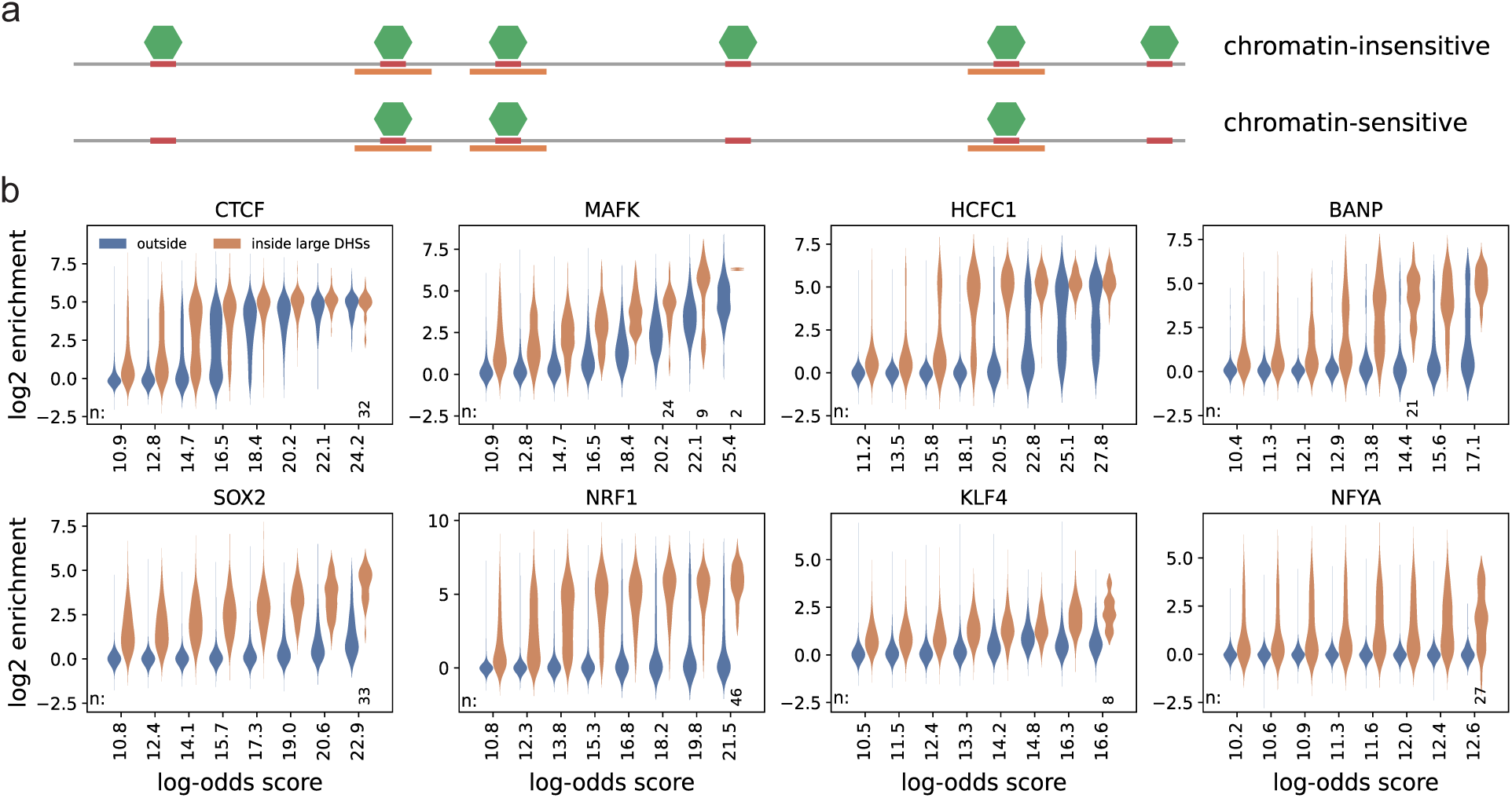
(a) Binding at motifs of a given strength (red rectangles) independent of chromatin context (e.g. large accessible regions, orange rectangles) indicates chromatin insensitivity, whereas a correlation between context and binding indicates chromatin sensitivity. Green hexagons represent TFs. (b) Binding versus log-odds score as in Figure 1c, but stratified by overlap with large DHSs. The number of motifs per group (n) are indicated if smaller than 50.

Our analysis has so far focused on binding behaviour at single motif instances. However, for several TFs, it has been shown that binding peaks often contain multiple motifs (e.g. Wang et al., 2012). As multiple motifs of the same TF could be regarded as an additional binding mode, similarly to a palindromic motif corresponding to two properly oriented binding motifs for the binding of a homo-dimer, it is therefore not a priori clear whether chromatin sensitivity should be defined via the binding behaviour at single motifs or also at clusters of motifs; however, estimating the relative affinity of clusters of motifs appears difficult as it may be a complicated function of the log-odds scores of each individual motif, their orientations and the inter-motif distances (Wang et al., 2012).

Additionally, even if relative affinities could be accurately estimated, a DHS-size-based approach would not allow to control for binding of other additional TFs in between the motifs. Due to these constraints, we here define chromatin sensitivity via the binding behaviour at single motif instances; it is accordingly important to ensure that the measured enrichments are not due to the presence of additional motifs of the same TF within a short genomic distance, whose contribution would be hard to distinguish due to the low resolution of ChIP-seq. We therefore reanalysed binding as a function of log-odds score and the presence of large DHSs using only motifs that do not have any other motifs (for the same TF) within +/- 200bps. This leads to the same qualitative results as when using all motifs (Supplementary Figure 6). While this solves the technical problem of multiple motifs in the current analysis, as we show below the problem of additional motifs close-by can also be circumvented using a deep-learning-based approach.

### Sequence-based quantification of chromatin sensitivity using Enformer

The genomic analyses so far have shown that chromatin sensitivity is continuous and generally depends on motif strength. However, their validity depends on a number of assumptions. First, using the amount of variability that cannot be explained by motif strength as an indicator of chromatin sensitivity assumes that the weight matrix models provide accurate estimates of relative affinity. Second, it assumes that motifs of different strengths roughly equally overlap with chromatin features that have an impact on binding. While our analysis shows that large DHSs are not equally distributed across log-odd bins, incorporating them into the analysis does not substantially change the inferred patterns of chromatin sensitivity. Nonetheless, this does not exclude that a more fine-tuned analysis, taking strength of DNAseI signal into account, will not reveal further dependencies.

Furthermore, while accessibility appears to be the strongest predictor of binding (Arvey et al. (2012) and also in our analysis (data not shown)), we cannot exclude that additional chromatin features may contribute to binding. Indeed, for some TFs, substantial binding variability remains at some motif strengths even outside of large DHSs (e.g. log-odds score bin 16.5 for CTCF or 25.1 for HCFC1 in Figure 2b), suggesting factors other than motif strength may be at play. Such predictive features may include, for example, epigenetic marks that dictate TF binding but that are not fully correlated with accessibility; binding of additional TFs whose binding does not create a substantial accessible site that can be measured with DNAseI, or creates one that is too small to be classified as a large DHS; and binding of factors in the immediate vicinity of the TF in question whose DNaseI signature cannot be separated from that of the target TF. Additionally, a TF could have a highly prevalent binding site that is available as part of a repetitive element, therefore providing multiple motif instances of various strengths co-occurring with relatively constant chromatin features.

We reasoned that if TF binding could be predicted from DNA sequence *in silico* at sufficiently high accuracy, the influence of the chromatin context could be determined with much higher confidence, as it would allow creating new motif/chromatin combinations that would control for the uneven genomic distributions described above. In addition, on the level of the DNA sequence, it should be more straightforward to separate the contributions of several TFs binding at close distance based on the location of their binding motifs, thus likely improving the capability to separate the effect of the motif of the TF in question from the effect of the chromatin, as encoded in the flanking sequence surrounding the motif. Finally, a sequence-based approach would override the need to choose the features (e.g. large DHSs) to be considered, since it would directly incorporate additional relevant sequence-encoded features that might potentially not be detected by an accessibility assay.

The deep learning tool Enformer (Avsec et al., 2021) has recently been shown to predict TF binding from sequence at impressively good accuracy. Enformer was trained on several thousand mouse and human datasets including CAGE, ChIP-seq for histone marks and TFs, DNaseI, and ATAC-seq, and predicts binding at a resolution of 128bps, which is comparable to the resolution of ChIP-seq. To investigate the feasibility of using Enformer to investigate chromatin sensitivity, we first assessed its prediction accuracy on our collection of ChIP-seq datasets by training it on each sample separately using transfer learning (Methods). Comparison of predicted to measured binding on the test set at the level of motifs (i.e. the subset of 128bp windows that overlap any predicted motif with log-odds score of at least 10 as in Figures 1c and 2b) revealed good prediction accuracy for many datasets (Figure 3a, Supplementary Figures 7-9), with the predictions clearly outperforming those of the log-odds score for all TFs (Figure 3a-b, Supplementary Figures 7 and 8). This suggests that either an improved assessment of motif strength or the information contained in the sequence flanking the motif (or a combination of both) leads to higher prediction accuracy. While predictive power generally decreases with lower motif complexity, good prediction accuracy is also observed for some TFs with lower complexity motifs (Supplementary Figure 10). This is particularly impressive for NFYA, whose low-complexity motif (Supplementary Figure 1) has very little predictive power, but for which Enformer predicts binding at good accuracy (Figure 3a-b).

**Figure 3.**
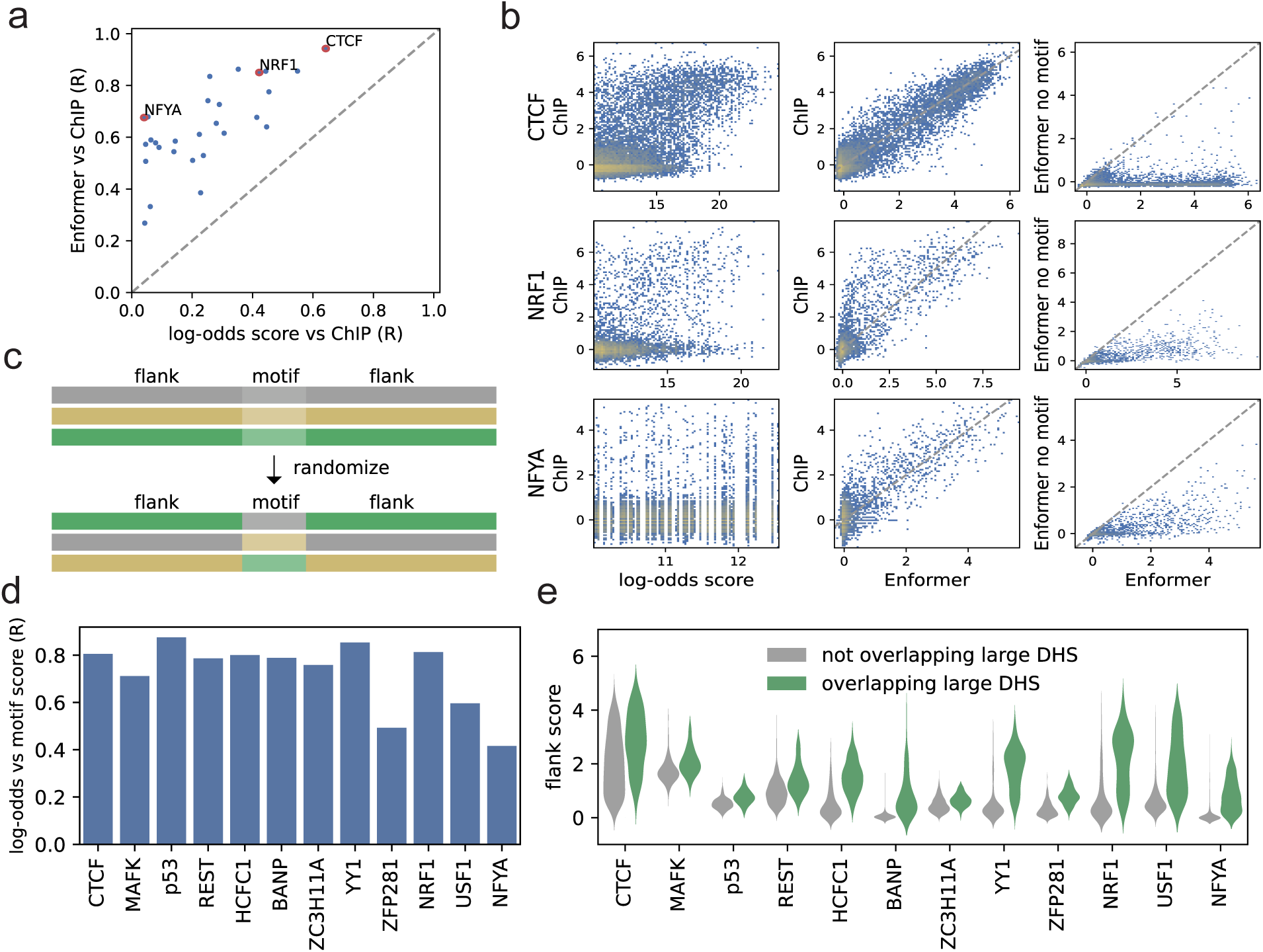
(a) Pearson correlation (R) of Enformer predictions or log-odds score vs measured ChIP enrichments (log2) on the level of motifs. (b) Predictive power of log-odds score versus Enformer for CTCF (top row), NRF1 (middle row) and NFYA (bottom row). ChIP-seq enrichment (log2) versus log-odds score (left) or Enformer predictions (middle) on the level of motifs and Enformer predictions in wt versus when all motifs have been mutated (right). (c) In-silico constructs are created by separating motifs from their endogenous flanking sequences (±500bps) and randomly recombining them. (d) Correlation between log-odds score and Enformer-based motif score for all TFs. (e) Distribution of flank scores stratified by whether the flanking sequence at its endogenous locus overlaps a large DHS. In d and e, TFs were sorted as in Figure 1b.

As mentioned above, the log-odds score analysis we employed heavily relies on the assumption that the available weight matrices for each TF are accurate and relevant.

Enformer allows testing the validity of the weight matrices used, by comparing binding predictions in wt to predictions where all motif sequences for a particular TF (log-odds score of at least 10) have been mutated genome-wide (Methods): A strong reduction of predicted binding signal upon mutation of these elements would indicate that the weight matrix was accurately inferred from the ChIP-seq data, whereas remaining signal would indicate that sequence elements other than the inferred motifs direct binding. Interestingly, comparison of predictions of wt and mutated sequences showed variable results, with samples from TFs with lower complexity motifs generally displaying a smaller reduction in signal upon mutation (Figure 3b right panel, Supplementary Figure 10). This may explain the low correlation between log-odds score and binding for some of the TFs in our dataset (Figure 1c, 2b and Supplementary Figures 2 and 5). For further analysis, we selected all samples that showed a good correlation between measured and predicted binding at motifs (R > 0.6), and where removal of the motif led to a large reduction in binding signal (mean relative loss in signal > 0.65, Supplementary Figure 10). This allowed us to use Enformer to investigate chromatin sensitivity for almost half of the samples in our dataset (12 out of 27).

As explained above, using Enformer to predict binding allows for the *in silico* construction of new motif/chromatin combinations that are not necessarily present in the genome, thereby directly assaying the contribution of chromatin, i.e. the features encoded by the DNA flanking the motif, to TF binding at different instances of the motif. We created such new motif/chromatin combination for each TF as follows. We randomly chose motifs in the genome as evenly as possible across the full range of log-odds scores as well as binding strengths (Methods). This ensures that, unlike in the genome (see Figure 1d), bound and unbound motifs as well as motifs of varying affinities are roughly equally represented. Additionally, in order to focus on motifs where the weight matrices represent the major binding mode, we only used genomic motifs where the predicted binding enrichment after mutation of the motif was below 2-fold over control (Methods, Supplementary Figures 7 and 8). For each such selected motif instance, we then split the motif from its surrounding 500bps sequence on each side, which we denote the ‘flank’, and created new motif-flank combinations by randomly combining motifs and flanking sequences (Figure 3c). As discussed above, this random pairing is intended to undo biases in the genomic distribution of motifs relative to chromatin features, as seen for large DHSs for a subset of TFs (Supplementary Figure 4). Importantly, if the flanking sequence contained additional predicted motifs for the investigated TF, these were mutated, in accordance with the framework described above that assesses chromatin sensitivity via binding behaviour at single motifs. As predictions with Enformer are computationally expensive, we devised a strategy to speed up predictions by predicting multiple instances within single 200kb stretches (Methods). This strategy also shows that while Enformer uses 200kb of input sequence, restricting the sequence context around a motif to *±*500bps leads to very similar predictions, justifying our flank definition (Methods, Supplementary Figure 11). Using this approach allowed us to query on average 630 different motif loci including their endogneous flanks per TF, such that each motif or flank appeared on average in 65 random motif-flank combination (Methods).

In order to be able to rank motif and flanking sequences according to the degree by which they promote binding, we defined a ‘motif score’ and a ‘flank score’, which represent the average predicted binding at all constructs with a given motif or flank, respectively.

Interestingly, except for NFYA and ZFP281, the motif score generally correlates well with the log-odds score (Figure 3d). For NFYA the low correlation may be due to the limited variability in log-odds scores within the investigated range of scores due to the low complexity of the motif, while for ZFP281, it is due to substantial differences at low log-odds scores (Supplementary Figure 12). The latter, albeit to a lesser extent, can also be observed for MAFK and YY1. Additionally, for some TFs, such as REST, p53, and HCFC1, there is a large spread in log-odds scores at low motif scores (Supplementary Figure 12). As Enformer has higher predictive power than the log-odds score, the motif score is likely a better estimate of relative affinity, with the difference being most substantial for weaker motifs. Flank scores of flanks that at their endogenous position overlap with large DHSs are on average higher than those that do not (Figure 3e), in line with the findings of the genomic analysis. However, the large variability in flank scores that is not explained by the presence of large DHS highlights the potential in assigning a quantitative value to the contribution of the surrounding sequence.

Similarly to the genomic analysis, the Enformer predictions allow us to investigate (predicted) binding as a function of motif strength. However, unlike in the genomic analysis, thanks to the pairing of each flank with many motifs of varying strength, the effect of motif strength on binding can be assessed for each flanking sequence separately (Figure 4a). In line with the genomic analysis, CTCF is predicted to be unbound at the weakest sites and bound at the strongest sites for the large majority of flanks. At low flank scores, CTCF requires strong motifs for binding, whereas at high flank scores, binding is already observed at lower motif scores, indicating that lower motif affinity can be compensated for by genomic context. This provides an explanation for the high binding variability observed at motifs of intermediate strength in the genomic analyses (Figure 1c and 2b). Importantly, unlike in the genomic analysis, Enformer predicts some variability in binding even at the strongest motifs, indicating stronger flank dependence and thus chromatin sensitivity. The fact that such variability was not noted in the genomic analysis again emphasizes that motifs are not randomly distributed in the genome and that the high log-odds score motifs are likely not equally represented in different chromatin contexts, thus masking their actual chromatin sensitivity.

**Figure 4.**
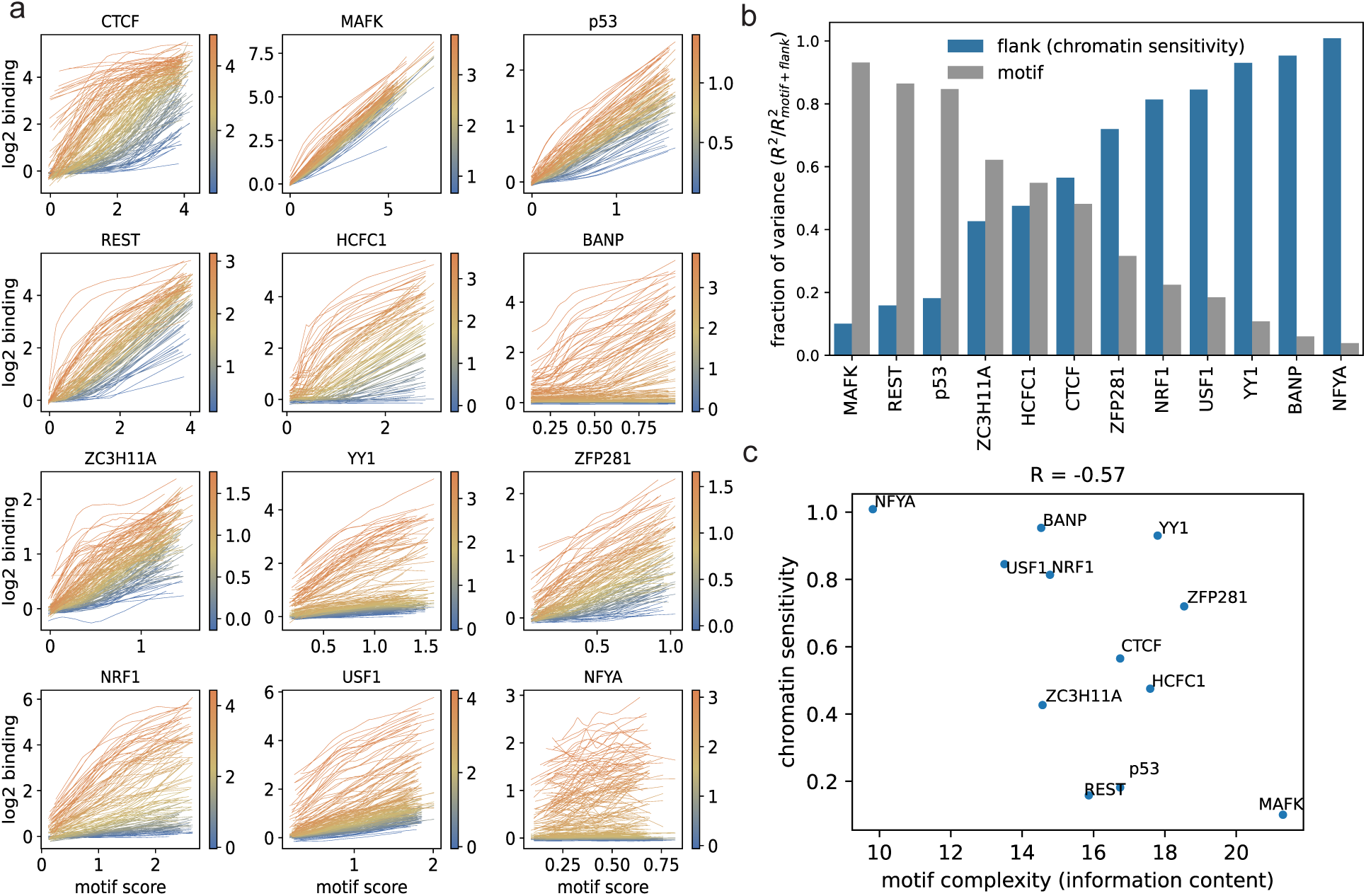
(a) Predicted binding versus motif score for different flanking sequences, coloured by flank score as indicated in the colorbars. Lines represent lowess regression lines (Methods). TFs are sorted according to the fraction of top bound motifs as in Figure 1b. To avoid overplotting, only a subset of flanks is shown for most TFs (Methods). (b) Fraction of variance explained by motif or flank score (R^2^), relative to an additive model that uses both motif and flank score to predict (R^2^_motif +flank_). TFs are sorted by increasing flank contribution. (c) Chromatin sensitivity (R^2^) versus motif complexity, as measured by the information content of the weight matrices. R indicates Pearson correlation coefficient.

For MAFK, predicted binding rises roughly linearly as a function of motif score, and exchanging a lower-scoring flank with a higher-scoring one leads to a constant and moderate rise in binding signal independent of motif score, indicating that MAFK displays moderate chromatin sensitivity irrespective of motif strength. The binding of NRF1 at high flank scores resembles that of CTCF at high flank scores, but unlike for CTCF, the lowest scoring flanks cannot be bound by NRF1 no matter how strong the motif, in line with the finding that NRF1 is chromatin-sensitive (Figure 1c and 2b). Similarly, HCFC1 and BANP (albeit based on only a small number of motifs in the test set for BANP, Supplementary Figures 8 and 9) do not bind their strongest motifs when embedded in lower-scoring flanking sequences, indicating they are chromatin-sensitive at their top motifs. Finally, NFYA binding strength is almost entirely a function of flank score, with a weak motif-score dependence at high flank scores, suggesting it is generally chromatin-sensitive and its chromatin sensitivity weakly increases with motif strength. It is important to note, however, that predicted NFYA binding never reaches enrichments as high as in wt (Figure 3b), likely due to the fact that in wt, additional NFYA motifs in the vicinity strongly contribute to binding (Wang et al., 2012). Indeed, when repeating the analysis without mutating additional NFYA motifs in the flanking sequence, the flank score strongly increases with the number of additional NFYA motifs (Supplementary Figure 13). An increase of flank score as a function of additional motifs in the flanking sequence can be observed for most of the TFs in our dataset (Supplementary Figure 13).

Taken together, the Enformer-based analysis indicates that binding is in general a complicated, non-linear function of both motif strength and genomic context. In general, the inferred patterns of chromatin sensitivity qualitatively agree with the genomic analysis. Nonetheless, the Enformer-based analysis provides a finer-tuned quantitative assessment for the characteristics of various motif and flanks, and therefore of overall TF binding behaviour.

### Chromatin sensitivity is a continuum across TFs

Both the genomic and Enformer-based analysis indicate that chromatin sensitivity is not binary but rather continuous and the degree of chromatin sensitivity is a function of motif strength. We wondered whether it would be possible to rank TFs by a ‘global’ measure that assesses chromatin sensitivity in general - across all motifs - rather than for motifs of specific strengths. While the binding curves in Figure 4a are generally non-linear, the sum of motif and flank scores correlate highly with Enformer predictions (Pearson correlations between 0.86 and 0.97), suggesting that the motif and flank scores can, at least as a good approximation, be treated as independent variables. In Figure 4b, we show the fraction of variance in Enformer predictions explained by either the flank score or motif score, relative to a model that uses the sum of flank and motif score: As the sum of motif and flank score shows high predictive power, high flank contribution implies low motif contributions and vice versa (Figure 4b). Thus, most of the information is contained in either one of the two variables. This naturally points to a new, sequence-based measure of a TF’s chromatin sensitivity: the fraction of variability in binding explained by the flanking sequence, relative to that explained by motif and flanking sequences combined. According to this logic, binding of a chromatin-sensitive TF is mostly determined by the flanking sequence around its motifs, whereas a chromatin-insensitive TF’s binding behaviour shows low flank dependence and high motif dependence.

Interestingly, flank dependence and thus chromatin sensitivity is a continuum across TFs (Figure 4b). On one end of the spectrum, the binding of MAFK is mostly determined by the strength of its motifs, and on the other end, the binding of NFYA appears almost entirely dependent on the flanking sequence. We observe a negative correlation between the complexity of a TF’s motif and its inferred chromatin sensitivity (Figure 4c). While it is unclear to what extent log-odds scores and thus information content can be directly compared between TFs (Ma et al., 2015), this suggests that TFs with higher potential binding affinity to their motifs tend to be less chromatin-sensitive. In particular, it suggests that TFs whose motifs have low information content - and therefore can appear by chance many more times in the genome - are more reliant on the chromatin context to direct their binding to specific sites. This observed continuum of chromatin sensitivity neither agrees with the bimodality in chromatin sensitivity predicted by a purely hierarchical model, nor with a purely cooperative model, which predicts that all TFs are chromatin-sensitive. It rather suggests that binding patterns are a result of a combination of hierarchical and cooperative mechanisms (see Discussion).

We wondered whether the results for the 12 TFs in mouse ESCs could be generalized to more TFs and cell types. To this end, we used all human ChIP-seq datasets which had been used to initially train Enformer (Avsec et al., 2021) and for which there was a binding motif available in the Jaspar database (Methods, Khan et al. (2018)). As in the case of the mouse ESC datasets, we selected datasets that showed a good prediction accuracy at the level of motifs (R > 0.6) as well as a strong reduction in predicted binding signal when mutating all predicted motifs (mean relative loss in signal > 0.8). The larger latter cut-off than for the mouse datasets was motivated by the fact that these datasets had not gone through the same manual curation, and had thus not been controlled for biases in the same stringent way. For TFs with more than five different samples available, we only used the five samples that showed the largest loss in binding upon mutation of the motif. This left us with 68 datasets for 30 different TFs. Similarly to the mouse dataset, Enformer predictions could be well approximated by the sum of motif and flank scores for all TFs (Pearson correlation between 0.80 and 0.97), and while the inferred motif score generally agrees well with the log-odds score (Supplementary Figure 14), they disagree for a subset of motifs with low log-odds scores, providing additional evidence that low log-odds scores may be imprecise (Supplementary Figure 15). This includes MAFK, in agreement with the observations in mouse ESCs (Supplementary Figure 12).

For these human datasets we observe a similar continuum of chromatin sensitivities as measured by the contribution of the flanking sequence to binding (Figure 5a), with a similar (albeit lower) negative correlation between information content of the motif and chromatin sensitivity (Figure 5b). As in mouse ESCs, the most chromatin-insensitive TFs (among all tested TFs) include MAFK and REST, and chromatin sensitivity increases for CTCF, USF1, and NRF1. This is also evident when separating predicted binding curves by flanking sequence (Figure 5c). While the most chromatin-sensitive TF of all tested TFs in mouse ESCs, NFYA, is not part of the human dataset, the TF NFYB, which is in the same complex as NFYA (Dolfini et al., 2012), is the most chromatin-sensitive TF among the human samples. Interestingly, similarly to MAFK, the remaining two members of the small MAF protein family, MAFF and MAFG (Katsuoka & Yamamoto, 2016), show a limited flank dependence and thus chromatin sensitivity (Figure 5a and c). It is possible that these TFs, due to their small size, can bind to linker regions between nucleosomes, making them less accessibility-dependent.

**Figure 5.**
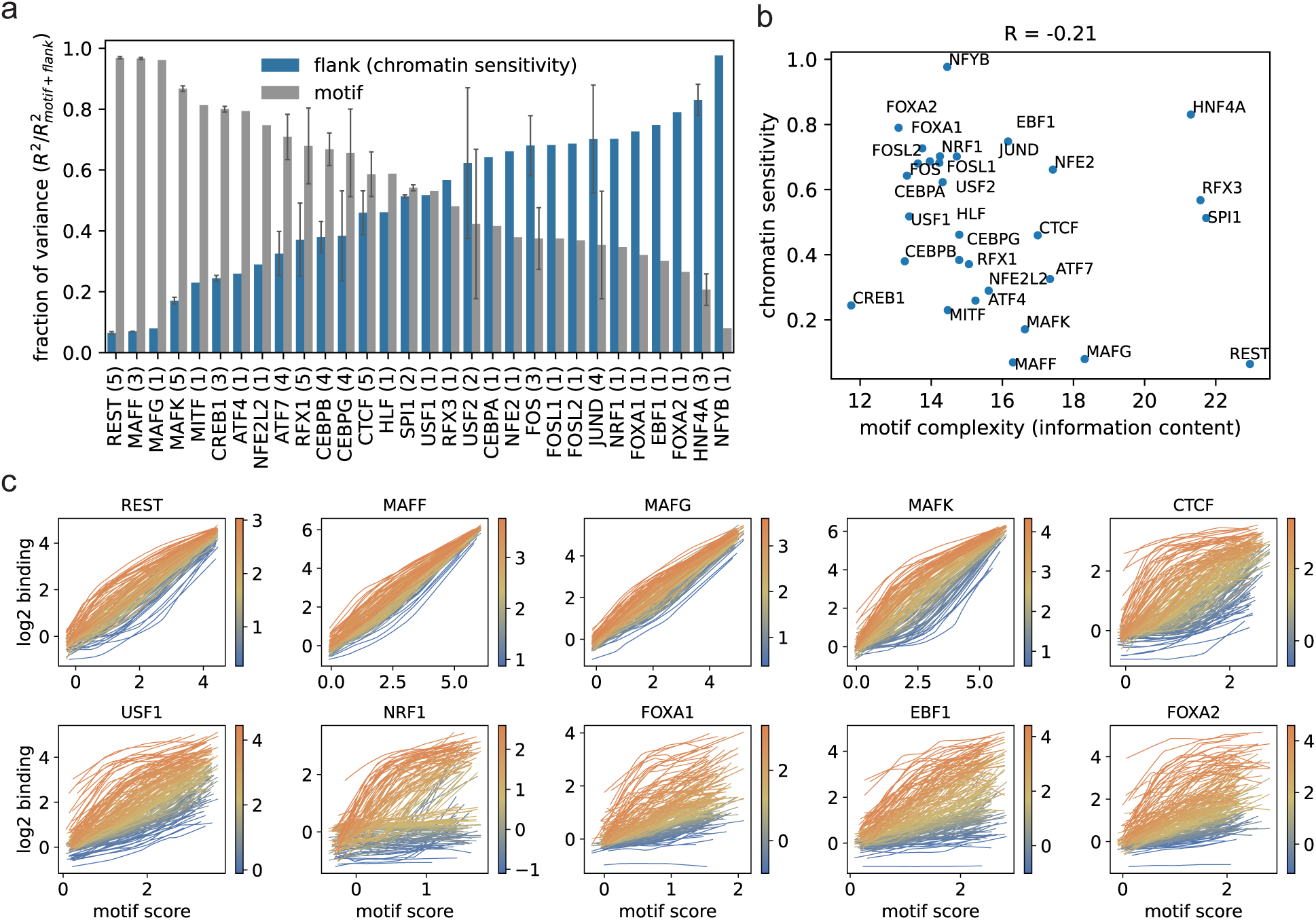
(a) Fraction of variance explained by motif or flanking sequence (R^2^), relative to an additive model that uses both motif and flanking sequence to predict (R^2^_motif +flank_) for human TFs. For TFs with multiple samples, average ± one standard deviation is shown. The number of samples is indicated in brackets. TFs are sorted by increasing chromatin sensitivity. (b) Chromatin sensitivity (R^2^) versus motif complexity, as measured by the information content of the weight matrices. R indicates Pearson correlation coefficient. (c) Predicted binding versus motif score for different flanking sequences for a selected set of TFs. Flanking sequences are colored by flank score as indicated in the colorbar. To avoid overplotting, only a subset of flanks is shown for every TF (Methods). For TFs with multiple samples, one randomly chosen sample is shown.

Among the more chromatin-sensitive TFs in our dataset are some TFs that have been classified as pioneer factors. For example, FOXA1 was the first TF to be named a pioneer factor based on its ability to initiate chromatin opening (Cirillo et al., 2002).

However, our analysis suggests that FOXA1 is strongly chromatin-sensitive (Figure 5a and c). This apparent chromatin sensitivity is due to the the presence of many low-scoring flanking sequences that are only very weakly bound even if combined with the strongest motifs (Figure 5c). In contrast, at sequences with high flank scores, binding increases strongly with motif score, indicating strong context-dependence; this is in line with the recent finding that FOXA1 is strongly affected by co-binding of other TFs (Xu et al., 2024). Some of the discrepancy may be explained by the fact that FOXA1 appears to be able to bind to inaccessible sites when they contain multiple motif instances (Hansen et al., 2022). This may either indicate that multiple motifs result in higher effective affinity, which allows FOXA1 to overcome chromatin, or it may indicate that at these loci, FOXA1 binds cooperatively with other factors that are also not able to open chromatin by themselves (see Discussion). For FOXA2, we find equally strong chromatin sensitivity and very similar binding behaviour as for FOXA1 (Figure 5a and c). This agrees with the finding that FOXA2 binds to only a subset of its predicted motifs in the genome and exhibits many cell-type specific binding sites (Donaghey et al., 2018). EBF1, a TF required for B cell differentiation, is also predicted to be strongly chromatin-sensitive (Figure 5a and c) and shows very similar binding curves to FOXA1 and FOXA2. This is in contrast to several studies that have shown that EBF1 is able to bind in closed chromatin (Boller et al., 2016; R. Li et al., 2018). Our finding suggests that even though the chromatin is closed, binding may only happen in places where there is cooperative binding with other factors which also cannot open the chromatin by themselves (see Discussion).

Taken together, the Enformer-based analysis consistently shows for both mouse and human samples that chromatin sensitivity is a continuum across TFs. This finding disagrees both with the bimodality predicted by a purely hierarchical model and with a purely cooperative model that implies that TFs are generally chromatin-sensitive. Our findings thus suggests that regulatory regions are activated by a combination of cooperative and hierarchical binding.

## Discussion

Using both genomic and deep-learning-based approaches, we have shown that a TF’s chromatin sensitivity is not solely a TF-intrinsic property, but also a function of the strength of the motif it encounters and thus locus-specific. There are likely additional TF-extrinsic contributing factors, the most obvious candidate being a TF’s expression levels (Blassberg et al., 2022; Gibson et al., 2024), which could influence its ability to bind to low-affinity sites that are otherwise bound in a chromatin-dependent manner. Here, our Enformer-based framework enables us to look across all motif strengths available in the genome and assign, for the specific cell types / expression levels of a TF, a ‘global’ measure of chromatin sensitivity, which assesses the relative contributions of the motif sequence and the flanking sequence to binding at all potential binding sites. It indicates that chromatin sensitivity is a continuum across many mouse and human TFs. The fact that this continuum spans a broad range of chromatin sensitivities seems to argue against a purely cooperative model of opening up of regulatory regions, as we also identify multiple TFs that are mostly chromatin-insensitive. Similarly, the existence of many TFs of intermediate chromatin sensitivity argues against a purely hierarchical model, which would predict a bimodal distribution of chromatin sensitivities. While this implies that regulatory regions are likely opened by a combination of both mechanisms, it does not rule out that for a given locus, only one of the two mechanisms is at work. In fact, which mechanism - cooperative, hierarchical, or a combination thereof - is dominant for a given locus depends on the particular combinations of TFs that bind it as well their chromatin sensitivities at the particular motif variants that are present. An interesting future line of work would be to investigate the prevalence of the two mechanism at specific regulatory regions that are essential to developmental or differentiation processes. Of note, while we believe that the inferred continuum of chromatin sensitivities is a robust feature, the exact degree of chromatin sensitivity of particular factors may require more in-depth investigations as they might be dependent on additional properties, e.g. the existence of monomeric versus dimeric motif variants.

While we see here that many TFs that have been defined as pioneers (based on their ability to bind in closed chromatin) actually display chromatin-sensitive behaviour, this is likely due to an important distinction: Some instances of binding in closed chromatin could be due to particularly strong motif instances or a particular (even if ‘closed’) chromatin environment. The latter could, for example, correspond to regions with binding motifs for two TFs, both of which cannot bind their motifs by themselves, leading to closed chromatin in the absence of either TF and to open chromatin only in the presence of both; in a cell type in which one of these TFs is present and the other is introduced, some previously-closed regulatory regions would open, leading to the classification of the latter as a pioneer factor, though in fact this is an example of cooperative binding. Therefore, while pioneering may describe the tendency to bind in previously closed regions - i.e. regions that might be very important for defining new lineages or differentiation processes - it is the comparison of where a TF binds to where it could potentially bind (taking into account motif strengths) that provides the information needed to judge whether binding is chromatin-dependent.

Despite their often impressive predictive power, deep-learning-based approaches have been criticised for their lack of interpretability. In recent years, contribution scores have gained popularity (Shrikumar et al., 2017). These measure the effect of single-nucleotide changes on predicted binding and can, in the context of TF binding data, be used to identify sequence motifs that have a strong impact on predictions. For a given locus, this may allow the identification of both a TF’s binding motif as well as additional motifs or sequence features that have an impact on binding. Our method provides an alternative approach to assess binding at a given locus by separating the contribution of a TF’s binding motif and the flanking sequence around the motif. The flank score we propose allows a ranking of flanking sequences in terms of how strongly (and how generally) they enhance binding, without the need to identify motifs for additional factors or determine their contribution as a function of their strength, relative spacing, and orientation. Nonetheless, flank scores could be combined with contribution scores or motif finding approaches to identify the elements which make particular flanks potent.

Additionally, flanking sequences could be exchanged between different TFs to assess to what degree high flank scores are TF-specific or generally enhance binding, which could potentially give insights into the evolution of regulatory elements.

We believe that our Enformer-based approach is the first method to quantitatively assess chromatin sensitivity that is not (or much less) biased by the unequal distribution of motif variants relative to chromatin features in the genome. To our knowledge, all previously proposed computational approaches rely on an assessment of binding or accessibility data (or a combination thereof) at genomic loci (Arora et al., 2023; Peng et al., 2024; Pop et al., 2023; Sherwood et al., 2014; Srivastava et al., 2021). In the method most comparable to the genomic analyses presented here, TFs are classified as accessibility-dependent or -independent based on whether or not incorporation of accessibility data leads to an improvement in the sequence-based prediction of ChIP-seq peaks (Pop et al., 2023). While similar in its basic idea to our analysis using large DHSs, this method only models binding at strong peaks. As we have shown, such a restriction results in an incomplete picture with respect to chromatin sensitivity. A different method, called PIQ, infers TF binding at predicted motifs from DNaseI profiles and uses the distribution of DNaseI signal around motifs to distinguish between predicted pioneering factors and accessibility-dependent ‘settler’ factors (Sherwood et al., 2014). While difficult to directly compare, we believe that PIQ is less controlled than our approach, as it is based solely on DNaseI data (and not on ChIP-seq data) and its predictions depend on the correct inference of TF binding profiles as well as a correct deconvolution of the DNaseI profiles of multiple close-by binding events. In contrast, our Enformer-based approach is directly based on binding data and separates binding of a TF from binding of additional TFs on the level of DNA sequence, which is likely more straightforward. The need to deconvolve DNAseI profiles of multiple TFs can be circumvented by measuring the chromatin state prior to induction of expression of the TF in question. Recently, deep-learning based approaches have been proposed to assess the relative contribution of sequence and chromatin features using such datasets (Arora et al., 2023; Durdu et al., 2024; Srivastava et al., 2021). While these likely represent the most controlled approaches, they rely on experimental setups where chromatin states can be measured independent of TF binding, and thus cannot be generally applied across many ChIP-seq datasets.

We have shown that for most TFs with low-complexity motifs, their mutation does not result in a strong loss of predicted binding signal. This may either indicate that ChIP-seq datasets for these TFs are of lower quality or suffer from biases, or that weight matrices sometimes do not accurately model a TF’s binding specificity. To test the latter, a direction of future investigation could be to use more complex motif definitions, for example via kmer-based models (Arvey et al., 2012; Ghandi et al., 2014) or directly via contribution scores, to more precisely define the set of motif variants of a given TF. Such extended motif sets may lead to a more substantial loss of predicted binding signal upon mutation. If so, individual motif variants of this set could be assigned motif scores using the approach described here and may allow an extension of our approach to a larger class of TFs.

Finally, a limitation of the Enformer-based approach is that for a substantial number of TFs, binding cannot be predicted at high enough accuracy to allow a quantification of chromatin sensitivity. While some of these cases may be due to low-quality ChIP-seq datasets, we expect future deep-learning methods to generally improve binding predictions and thus more datasets should become accessible to such an analysis.

## Acknowledgements

We thank all members of the D.S. laboratory for their critical input on the study, Michael Stadler and Ana Petracovici for their critical input on the manuscript. D.S. acknowledges support from the Novartis Research Foundation, the Swiss National Science Foundation (310030_212716 and 310030B_176394 to D.S.) and the European Research Council under the European Union’s (EU) Horizon 2020 research and innovation programme grant agreements (DNAaccess884664)

## Author contributions

D.S., L.B. and N.G. conceived and designed the study, L.B. performed the analyses, and L.B. and N.G. wrote the paper.

## Methods

### Genomic analysis

Analyses were performed using the mm10 mouse genome assembly as well as the hg38 human genome assembly. Promoters for mm10 (one per gene based on maximum PolII signal) were defined as described in Grand et al. (2021). Mouse ChIP-seq datasets were processed and binding quantified at predicted sites as described in Ginno et al. (2020). While the initial list of ChIP-seq datasets contained more than one replicate for many of the TFs, we selected the one with the highest enrichment (after making sure that replicates showed consistent enrichments), defined as the median log2 enrichment of the top 100 sites. For a summary of all data-sets used, see Table 1.

For violin plots of binding enrichments versus log-odds score (Figure 1c and 2b and Supplementary Figures 2, 5 and 6), motifs for each TF were binned by log-odds score as follows: the motifs with the 250 highest log-odds scores were placed into the top bin and the remaining motifs were binned into 7 bins of equal width down to the lowest log-odds score (log-odds score of 10). While this resulted in 8 bins for most TFs, for a small number of TFs, certain bins contained no motifs, leading to a smaller number of bins.

DNaseI data for mouse ESCs (2 replicates) was downloaded from GSE67867 (Domcke et al., 2015), adaptors were trimmed using cutadapt (Martin, 2011) with parameters ‘-a GATCGGAAGAGC-m 15 –overlap=1’ and peaks were called using Macs2 (Zhang et al., 2008) with parameters ‘–nomodel –shift −25 –extsize 50 –keep-dup all’. The narrow peaks of the two replicates were merged and only those merged regions were retained that overlapped with peaks of both single replicates. This resulted in 184’664 peaks (DHSs).

### Enformer-based analysis

Enformer (Avsec et al., 2021) was downloaded from https://github.com/google-deepmind/deepmind-research/tree/master/enformer and a re-implementation for transfer learning from https://github.com/lucidrains/enformer-pytorch. Log2 binding signal was defined in mouse as the log2 enrichment over control as used for the genomic analysis. To calculate enrichments, each 128bp window was extended to 201bp centered on the mid-points of the 128bp windows to increased read counts per window. In human, log2 binding signal was defined as the log2 signal of the data used for training Enformer (Avsec et al., 2021) (which is based on the data from (Kelley, 2020)) minus the average log2 signal over all 128bp windows. Only samples that were untreated (according to the description column of the annotation table provided) were used.

Transfer learning was performed one sample at a time, retraining the last layer of the network. As this is a linear layer, we simply performed a linear (ridge) regression on the input values to this layer using scikit-learn (Pedregosa et al., 2011) with *α* = 0.01. Due to memory limitations and as the number of parameters to fit (3073) is much smaller than the total number of training samples (7323 *·* 896), we only used 25 percent of the training data to fit the model.

To speed up Enformer predictions, we made use of the fact that Enformer predicts binding simultaneously for all 128bp windows of a more than 100kb long input sequence: we placed motifs combined with their flanking sequences at a sufficiently large distance (5kb) into the same input sequences, allowing a simultaneous read-out of predicted binding at multiple motif-flank combinations.

Binding sites were predicted genome-wide using either the MOODS (Korhonen et al., 2009) Python package or the R package Biostrings (Pagès et al., 2024), using the identified weight matrices for all TFs in mouse and the weight matrices of Jaspar 2018 (Khan et al., 2018; Tan, 2017) for human. Motifs were mutated by setting the corresponding nucleotides within each motif to Ns, which correspond to (0, 0, 0, 0) when used as input to Enformer. Mean relative loss in signal after mutation was measured as the average of (*x_wt_ − x_mut_*)*/x_wt_*, where *x_wt_* is the predicted log2 signal in wt and *x_mut_* the predicted signal after mutation of the motif. The average was calculated over motifs with *x_wt_ ≥* 1. For mouse, we used a cut-off of 0.65 on the mean relative loss in signal whereas for the human samples a cut-off of 0.8.

For the creation of *in-silico* motif-flank constructs, motifs were subsampled from an equal-width grid of log-odds score and binding, with 8 bins for the log-odds score and 3 bins for (log2) binding values. For each bin 1000*/*(3 *·* 8) motifs were randomly sampled. In cases where a bin contained fewer motifs, all motifs were selected. Additionally, although the samples used for the Enformer-based analysis displayed a large loss in signal upon mutation of the motif, they sometimes still contained a subset of motifs with substantial log2 enrichment after mutation of the motif (log2 enrichment > 1). In order not to bias the analysis, these motifs were excluded from the subsampling procedure.

Lowess regression lines in Figures 4a and 5c were calculated using the Python package statsmodels (Seabold & Perktold, 2010) with *frac* = 0.5. To maximize the number of displayed flanks without too much overplotting, flanks were sorted by flank scores and in Figure 4a only every 3rd flank was plotted for USF1, YY1 and NFYA, every 5th flanks for the remaining TFs, except for BANP where every flank was shown. For all human samples, every 5th flanks is shown for each TF in Figure 5c.

Data analysis was performed in R and Python. All plots were made using the Python packages matplotlib (Hunter, 2007), seaborn (Waskom, 2021) and pandas (https://doi.org/10.5281/zenodo.3509134). Sequence logos were generated using the Python package Logomaker (Tareen & Kinney, 2020).

---

**Supplementary Figure 1.**
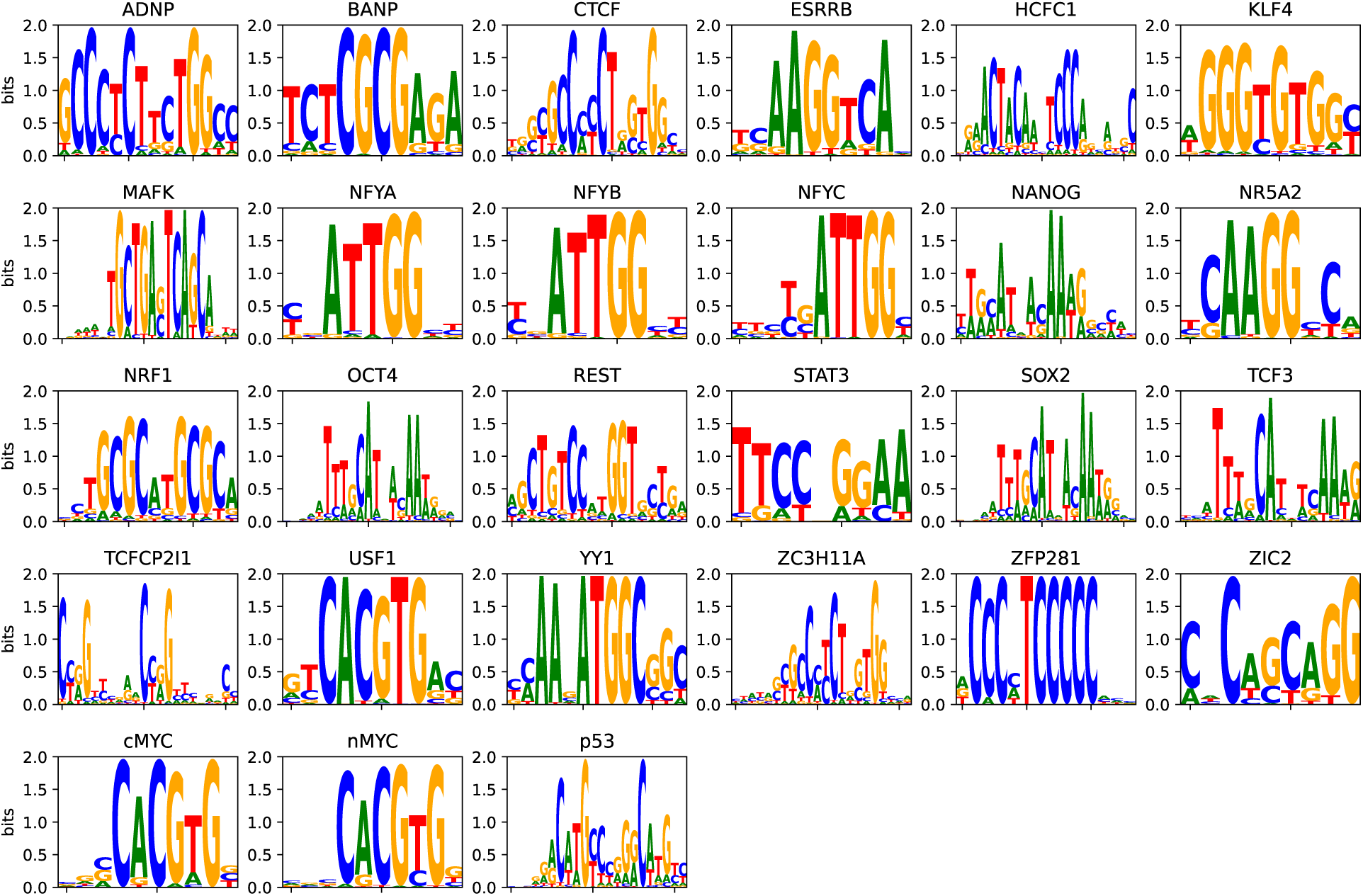
Weight matrices for all 27 TFs as inferred from the top 500 peaks.

**Supplementary Figure 2.**
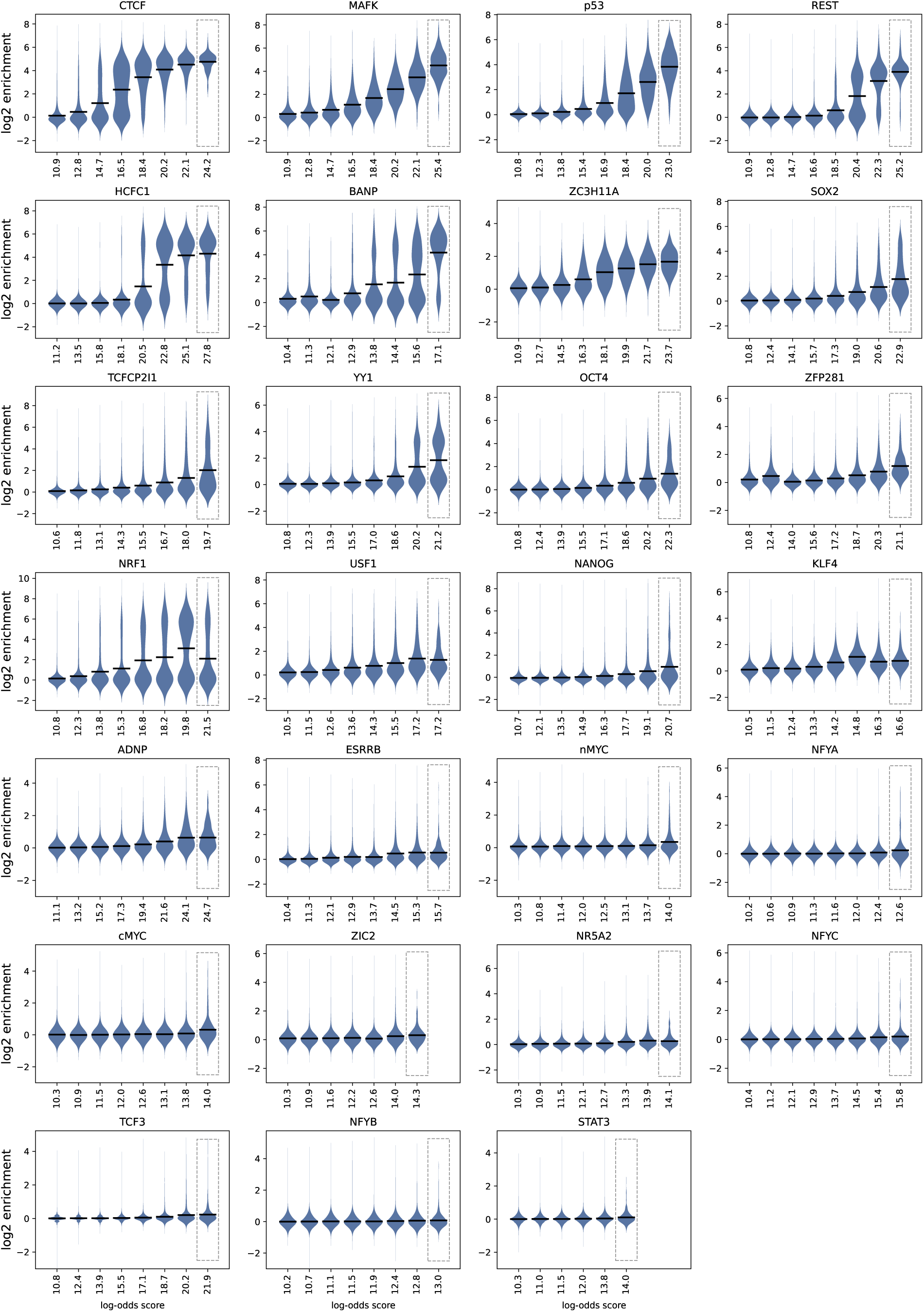
Binding enrichments versus log-odds score as in Figure 1c for all 27 TFs.

**Supplementary Figure 3.**
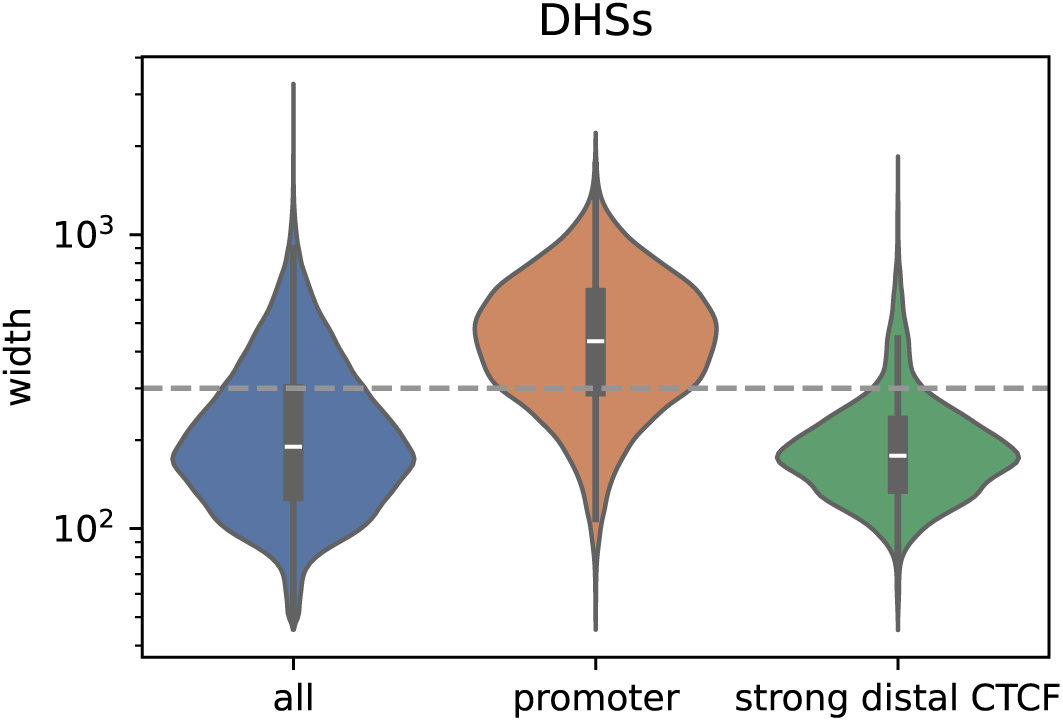
The width of DHSs in the genome (’all’), at promoters and at distal CTCF sites that are at least 4-fold bound. The dashed line marks the cut-offs to define large DHSs (300 nts).

**Supplementary Figure 4.**
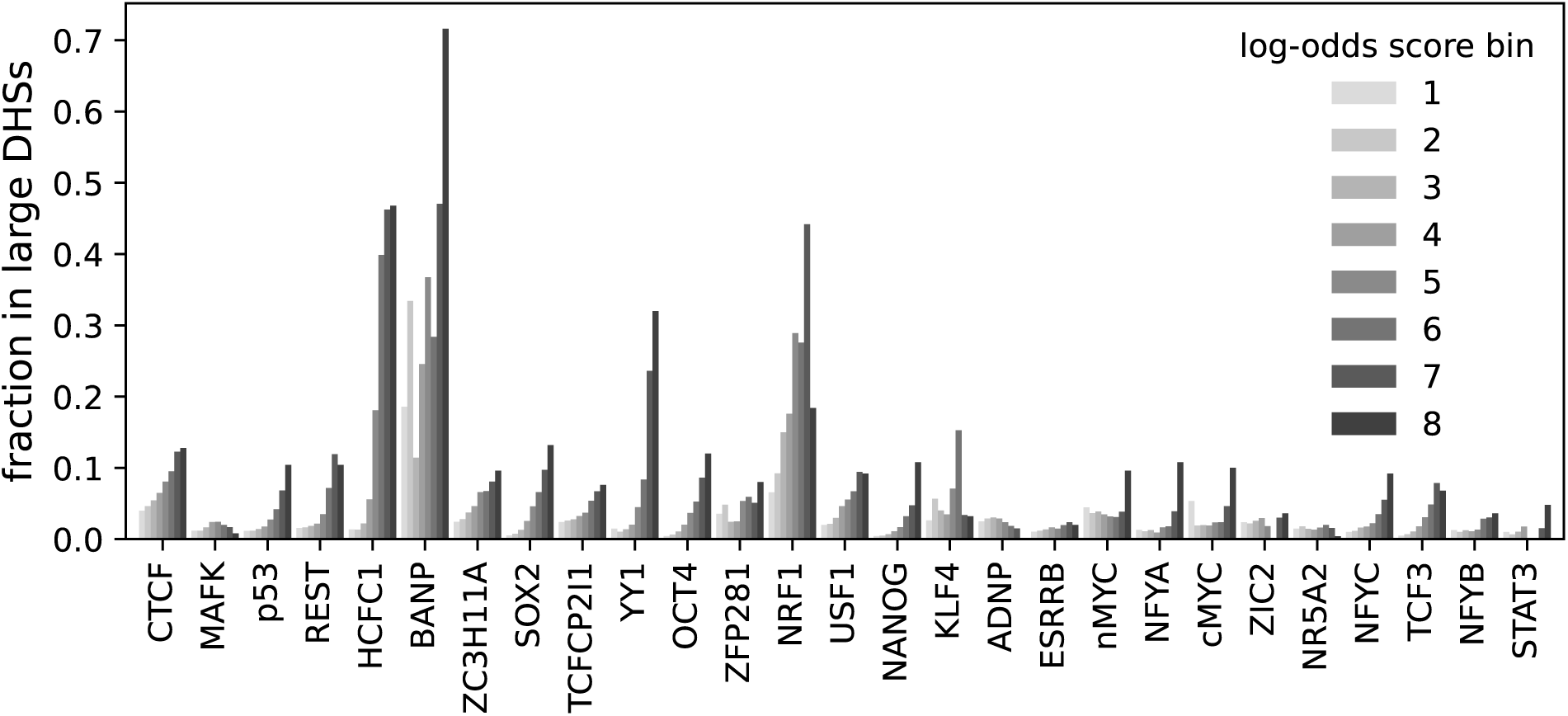
(d) Fraction of motifs overlapping large DHSs in each log-odds score bin (1: lowest, 8: highest). TFs were sorted as in Figure 1b.

**Supplementary Figure 5.**
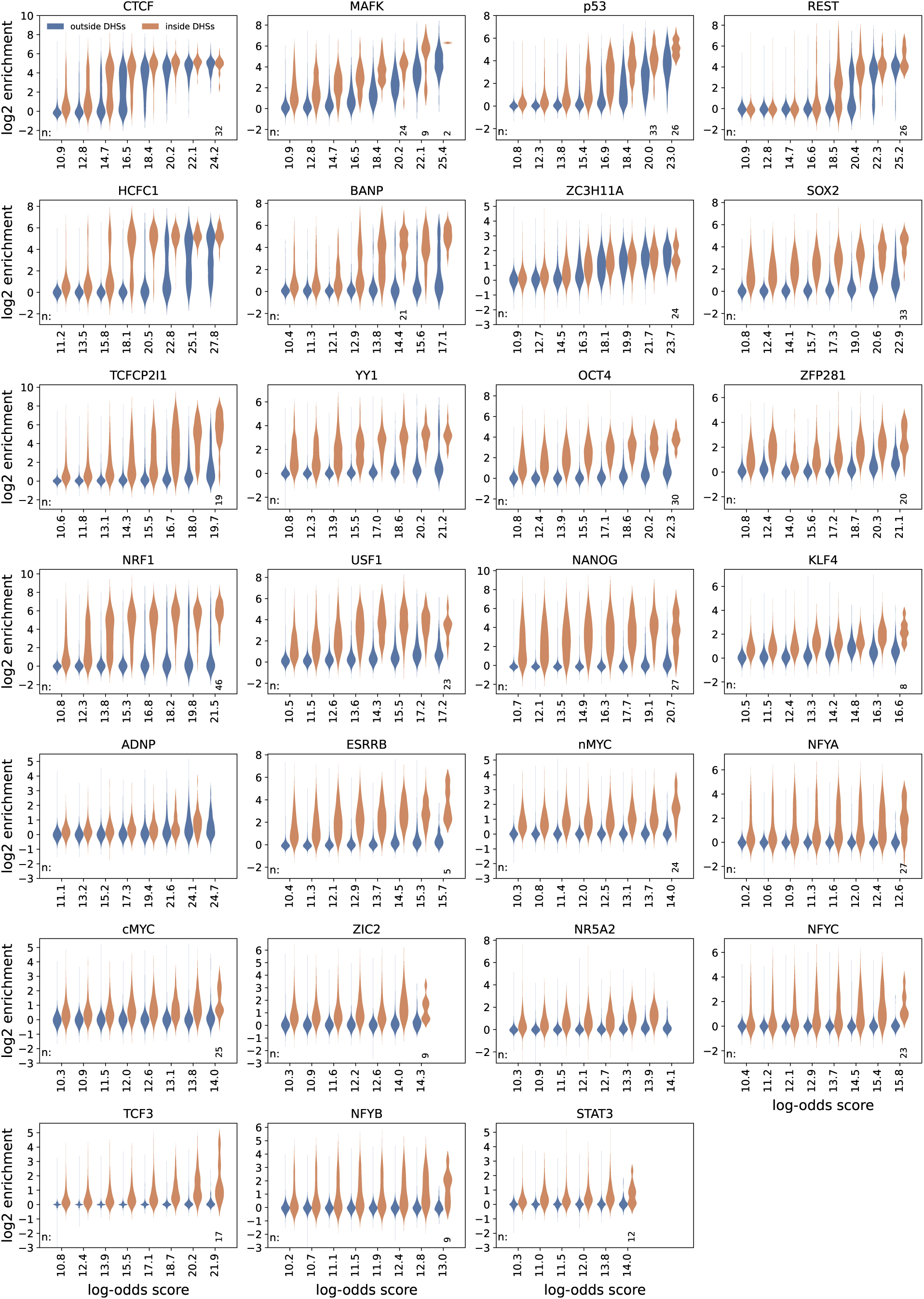
Binding versus log-odds score stratified by large DHSs as in Figure 2b for all datasets.

**Supplementary Figure 6.**
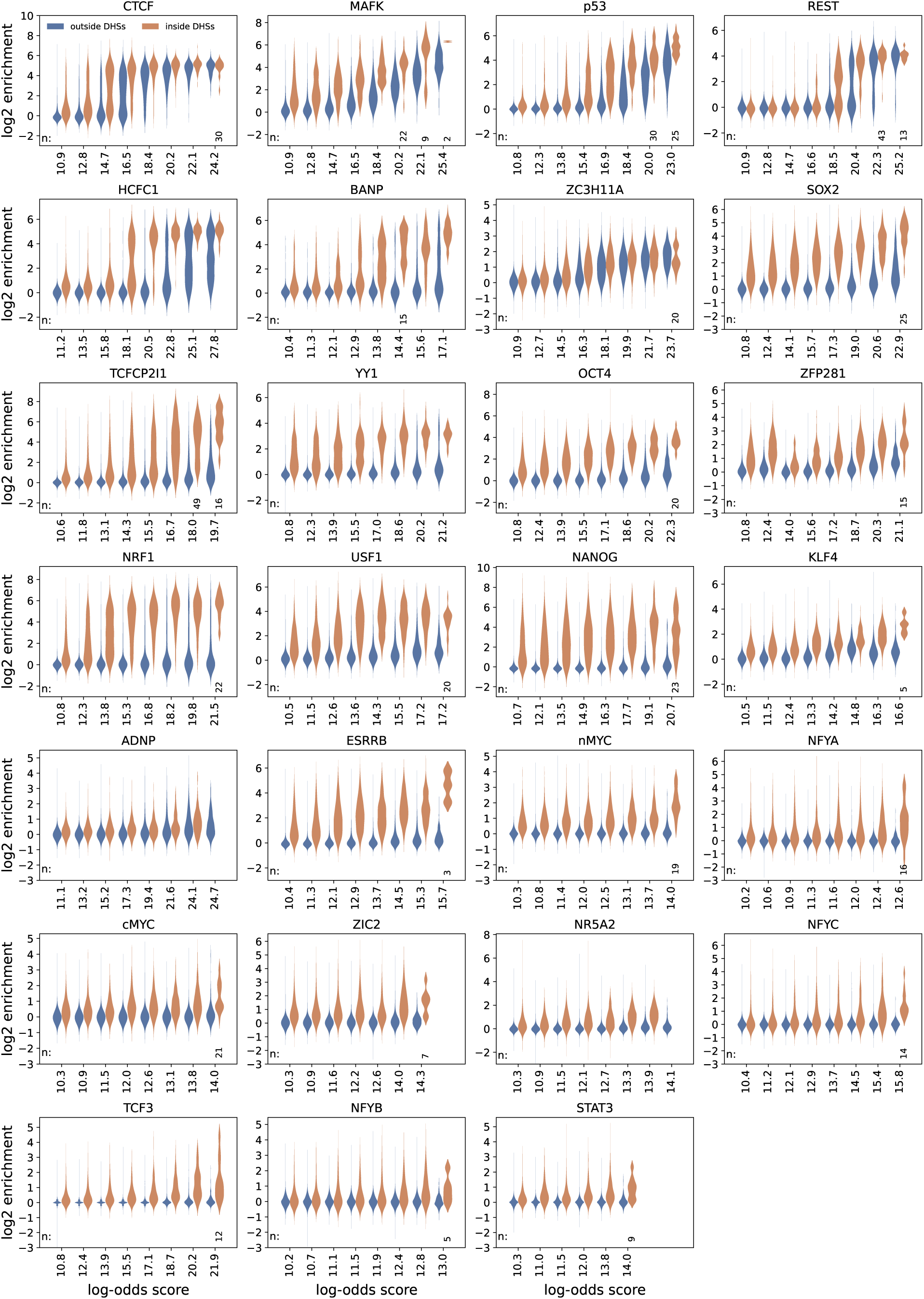
As Supplementary Figure 5 for motifs with no other motif (for same TF) within 200 bps.

**Supplementary Figure 7.**
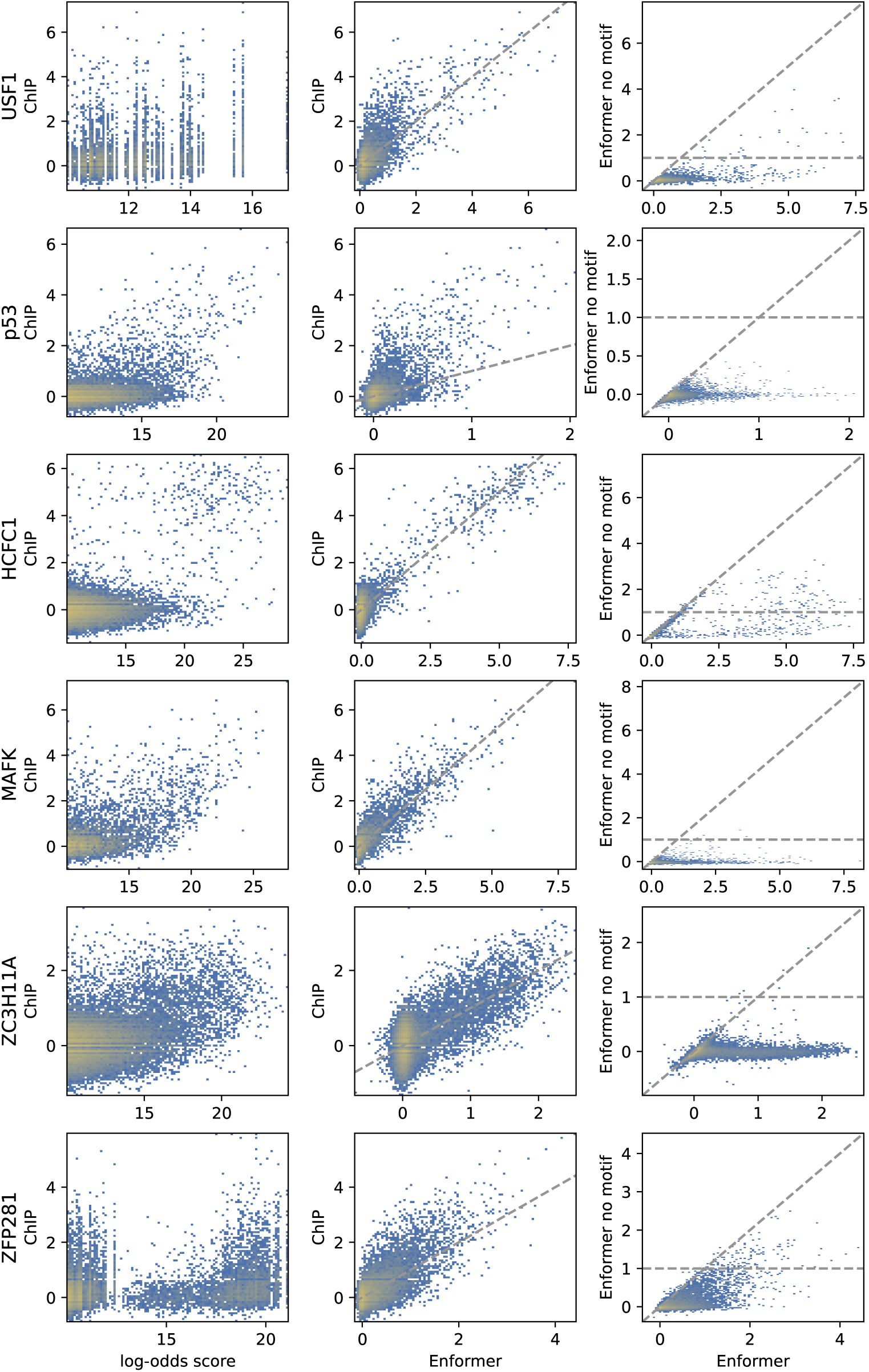
As Figure 3b, but for all TFs. For p53 and to a lesser extent ZFP281, predicted binding is generally smaller than measured binding (middle panel), but nonetheless roughly linearly related and thus should allow relative quantification of binding as a function of genomic context.

**Supplementary Figure 8.**
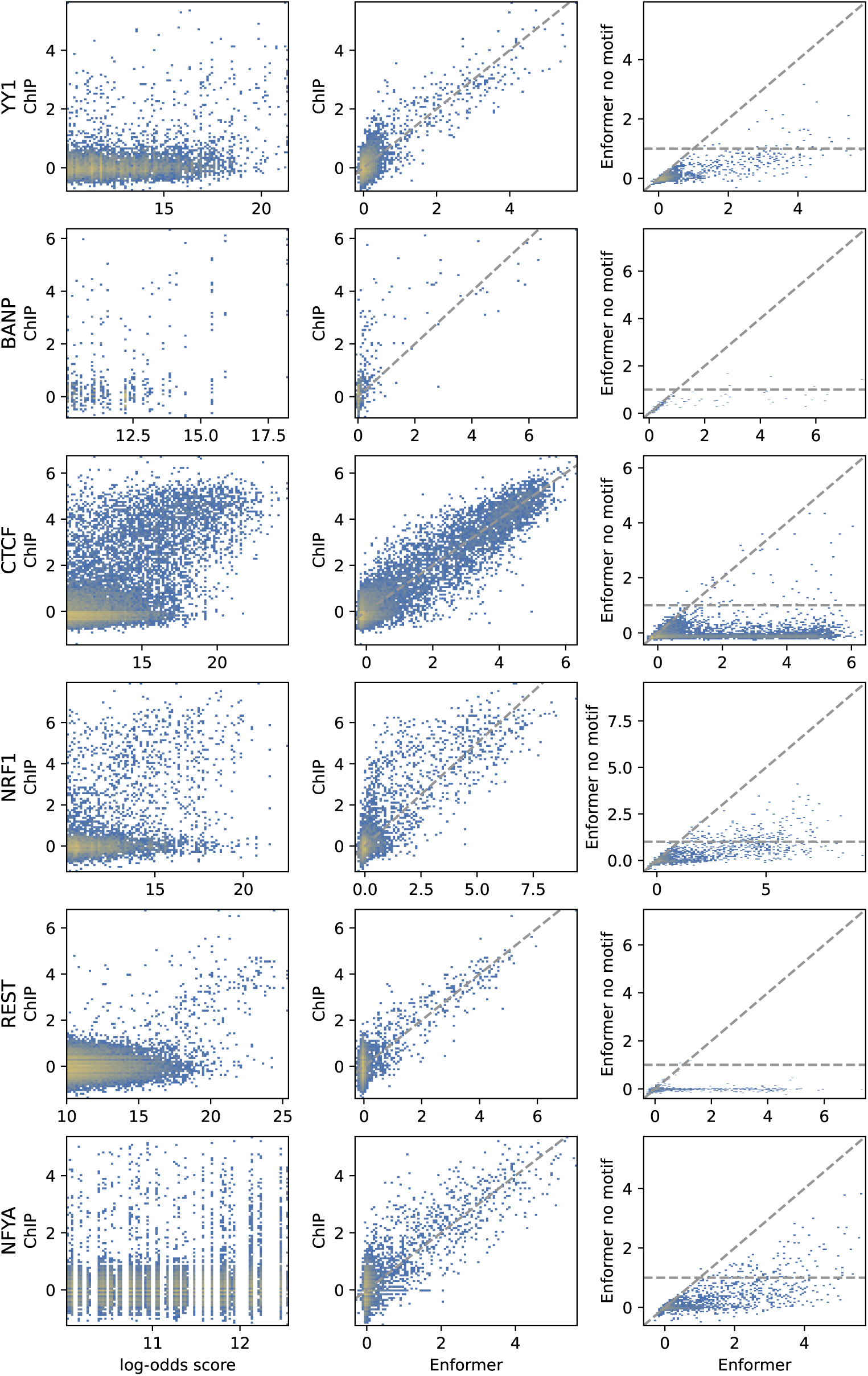
Continuation of Supplementary Figure 7. For BANP, due to its low number of target sites, the number of bound motifs in the test set is small. The good predictive power for BANP can be more clearly seen in Supplementary Figure 9.

**Supplementary Figure 9.**
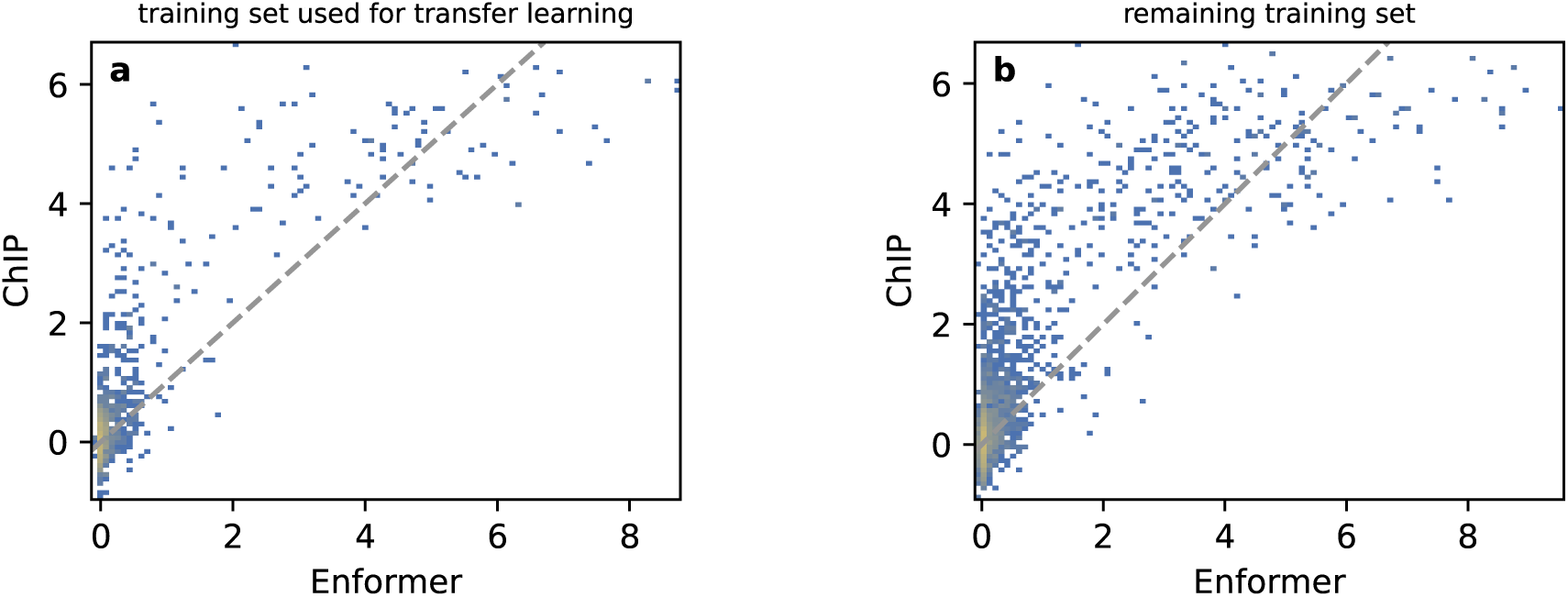
Measured (ChIP) versus predicted binding (Enformer) of BANP at all motifs overlapping the subset of the Enformer training set used for transfer learning (a) or the remaining regions of the Enformer training set, which were not used for transfer learning (b). For details, see Methods.

**Supplementary Figure 10.**
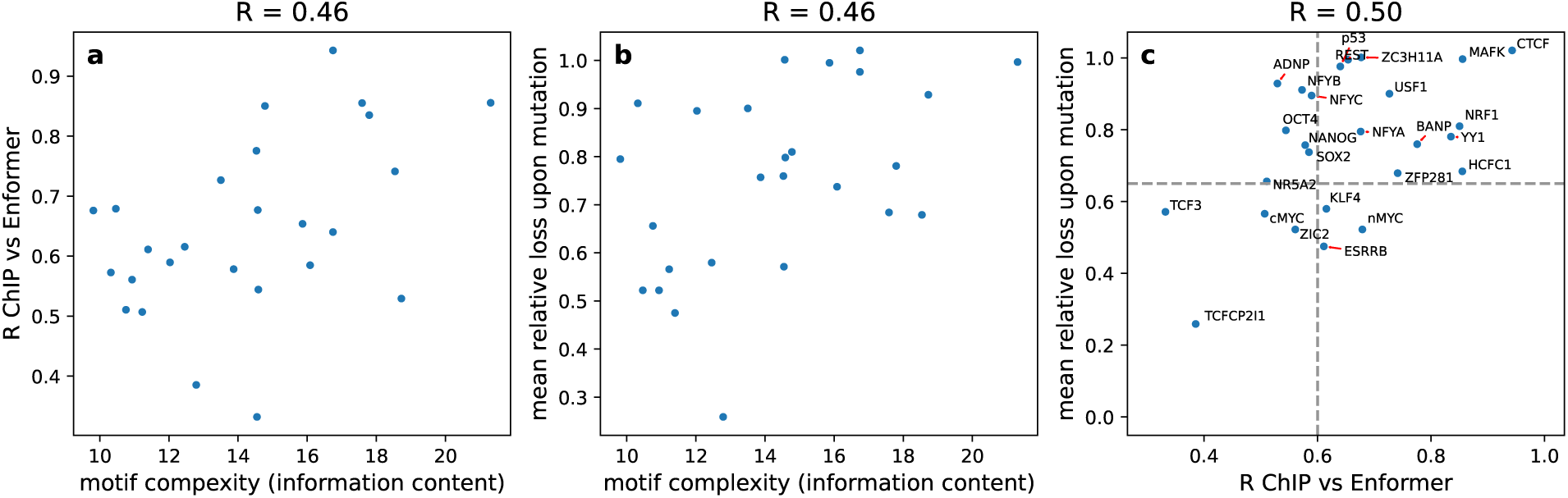
**a)** Pearson correlation (R) between measured and predicted binding at motifs versus the complexity of the motif, as measured by the information content of the weight matrices. b) Loss of predicted binding upon mutation of the motif versus information content of the weight matrix. Values close to one indicate that on average the signal is reduced to base-line upon mutation whereas values close to 0 indicate no change (Methods). c) Loss of predicted binding upon mutation of the motif versus Pearson correlation between measured and predicted binding at motifs. Dashed lines indicate cut-offs used for sample selection.

**Supplementary Figure 11.**
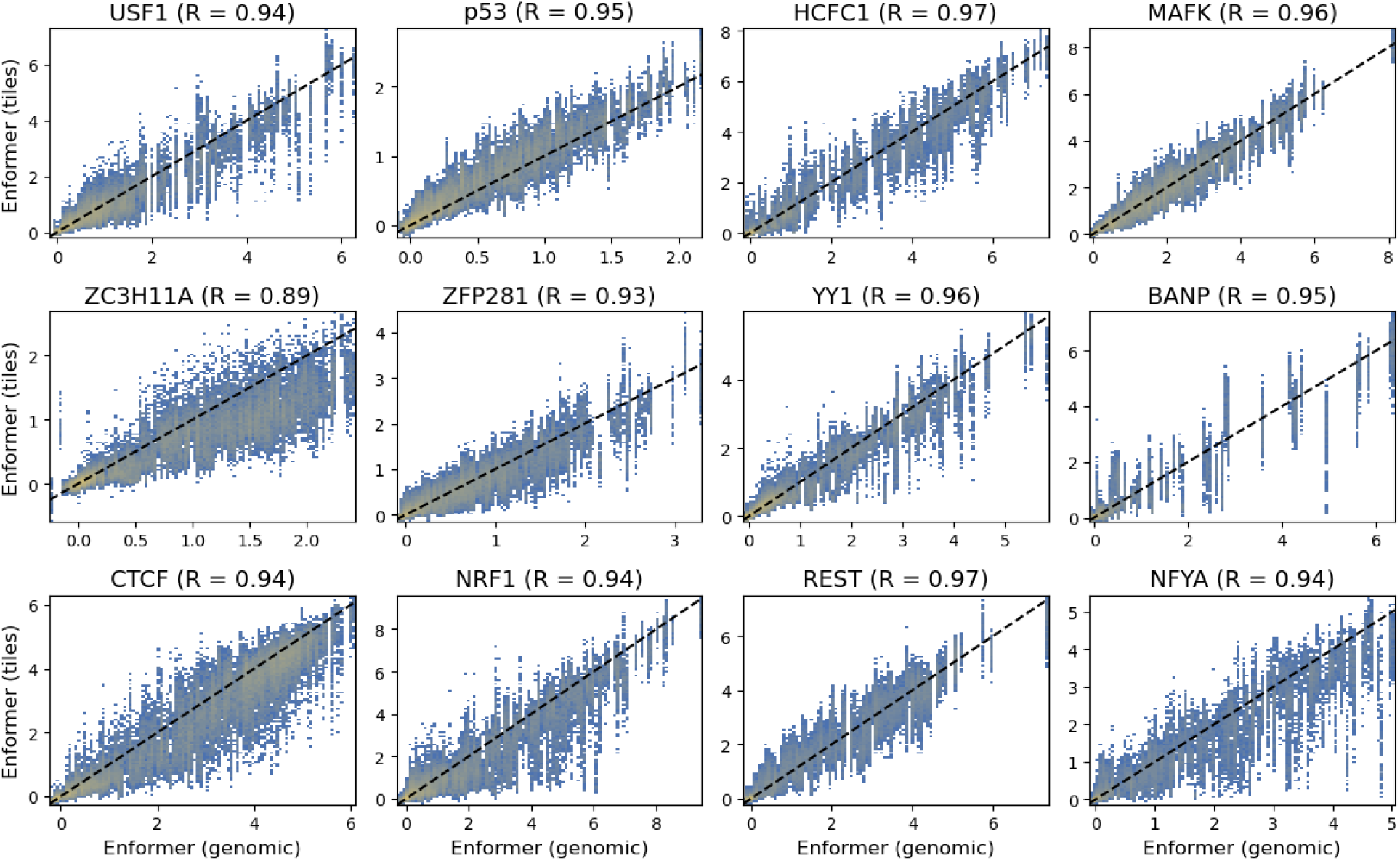
Enformer predictions at endogenous motifs (genomic) versus binding predictions when placing the same motifs ±500 bps of flanking sequence at 5kb distance along the genome (tiles, see Methods). Each motif (including its endogenous flanking sequence) was inserted multiple times, resulting in vertical lines in the plots. The comparison consists of all motifs that were used to create random motif-flank pairings (cf Figure 3c). While predictions for ZC3H11A are reduced when motifs are inserted along the genome, there is still a tight and roughly linear relationship, which should be sufficient for investigating relative binding strength as a function of different sequence contexts.

**Supplementary Figure 12.**
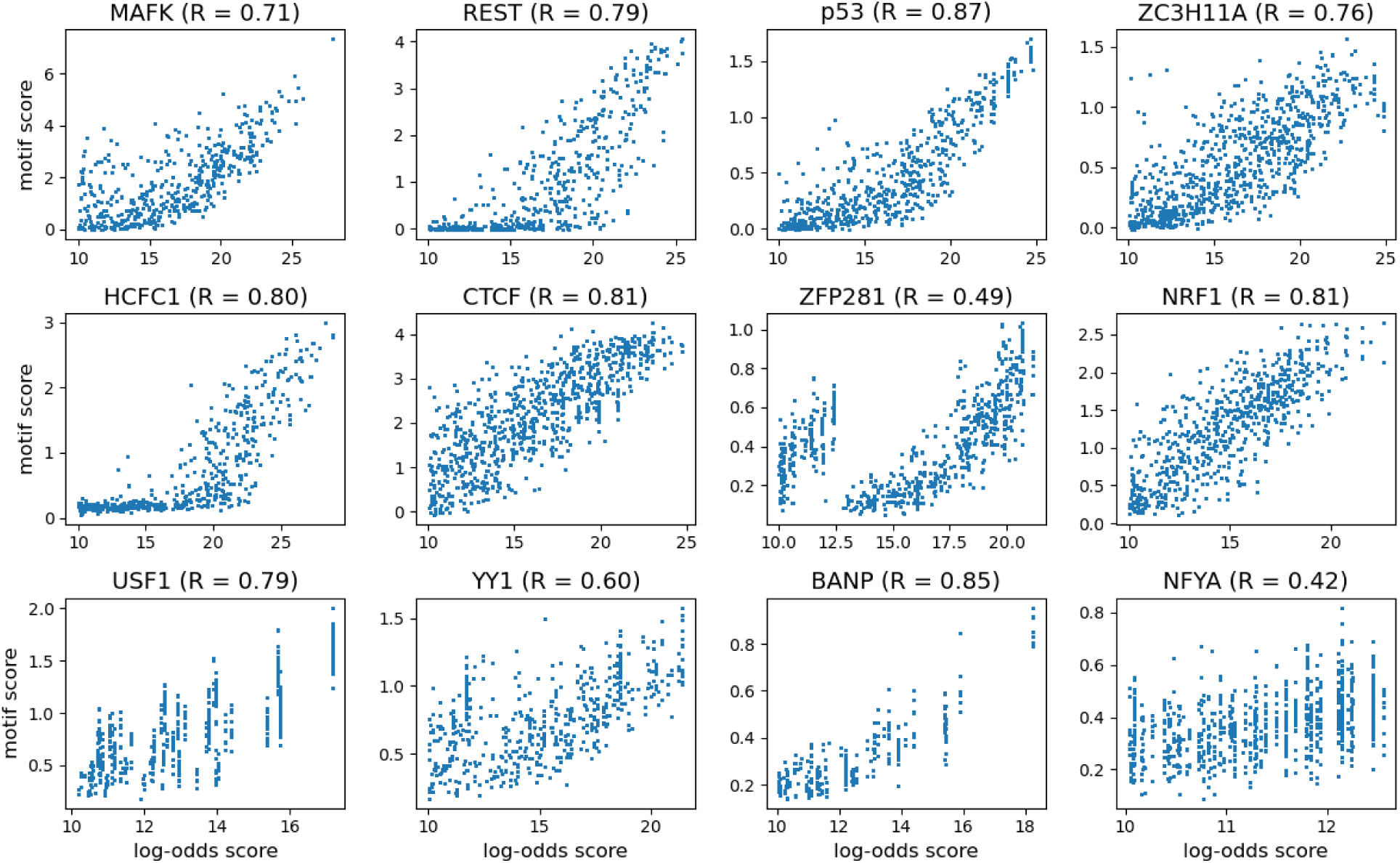
Log-odds score versus Enformer-based motif score. R indicates Pearson correlation.

**Supplementary Figure 13.**
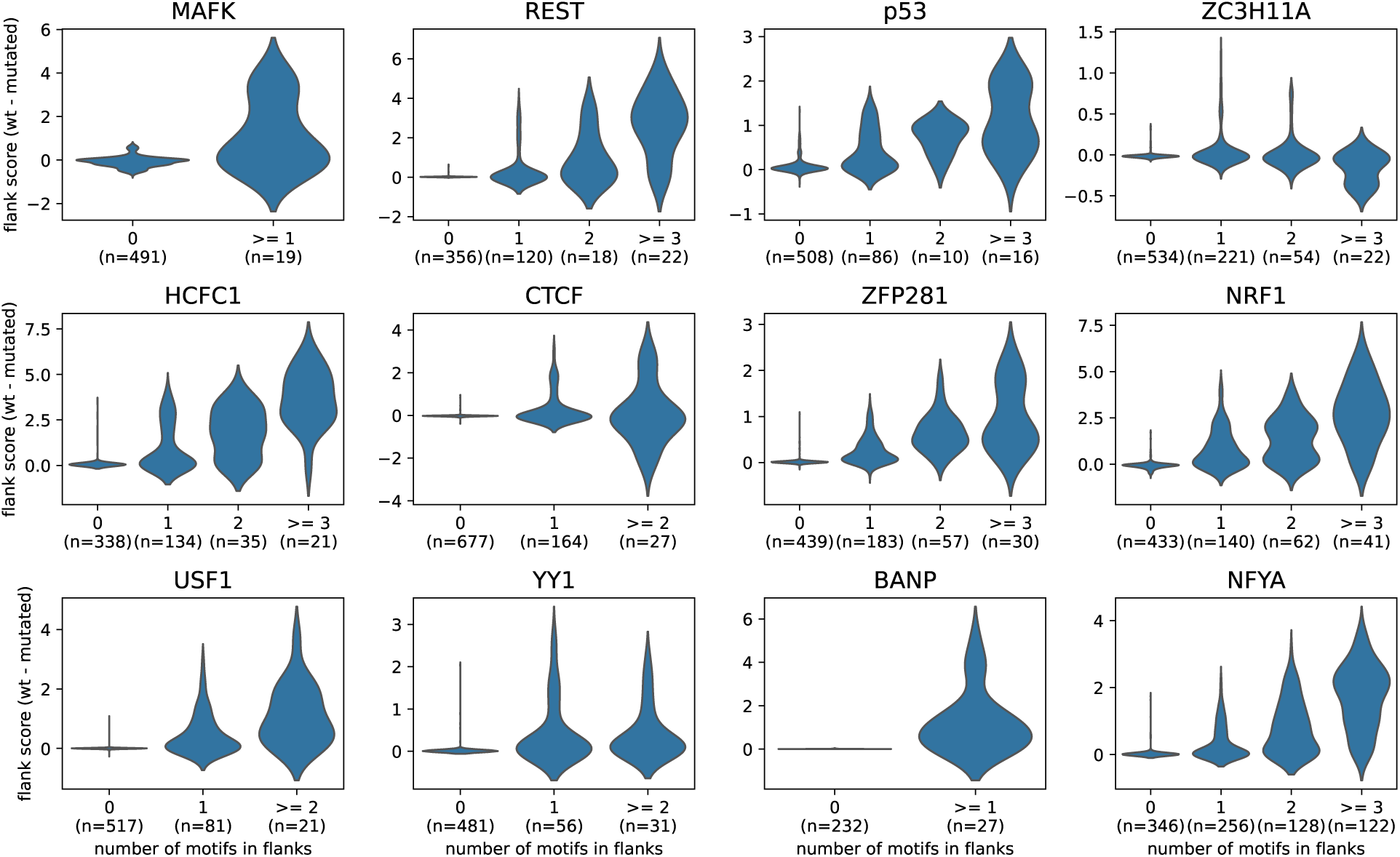
Change in flank score as a function of the number of additional motifs when using wt flanking sequences compared to flanking sequences with additional motifs mutated. The small variation observed at 0 additional motifs is due to the strategy used to speed up predictions where motif-flank constructs are placed along the same input sequence at sufficiently large distance (Methods): mutating motifs in a flank can have a small influence on predictions for other constructs not containing this flank.

**Supplementary Figure 14.**
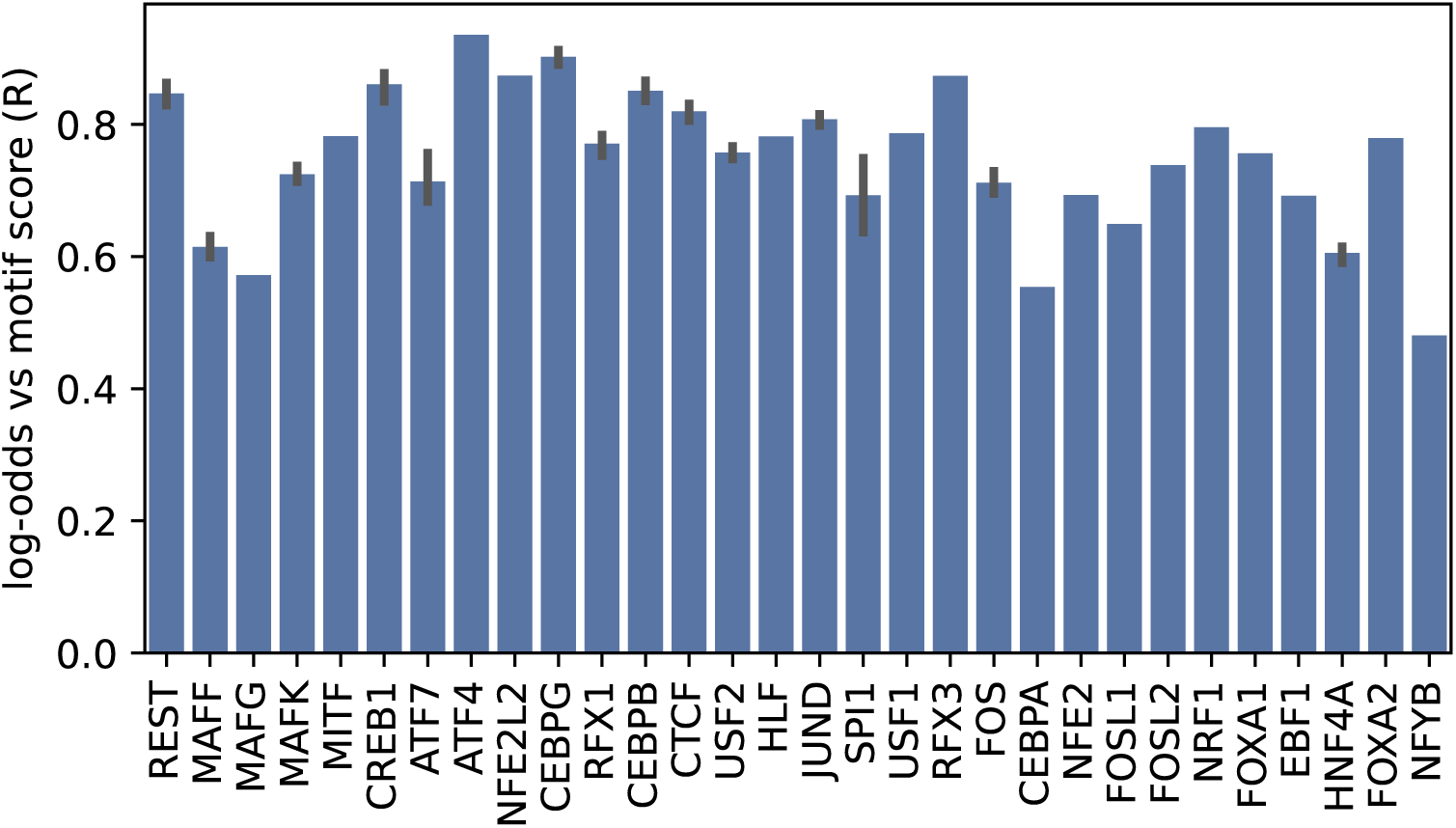
Pearson correlation between motif score and log-odds score for all human samples. For TFs with multiple samples, average ± one standard deviation is shown. TFs are sorted by increasing chromatin sensitivity.

**Supplementary Figure 15.**
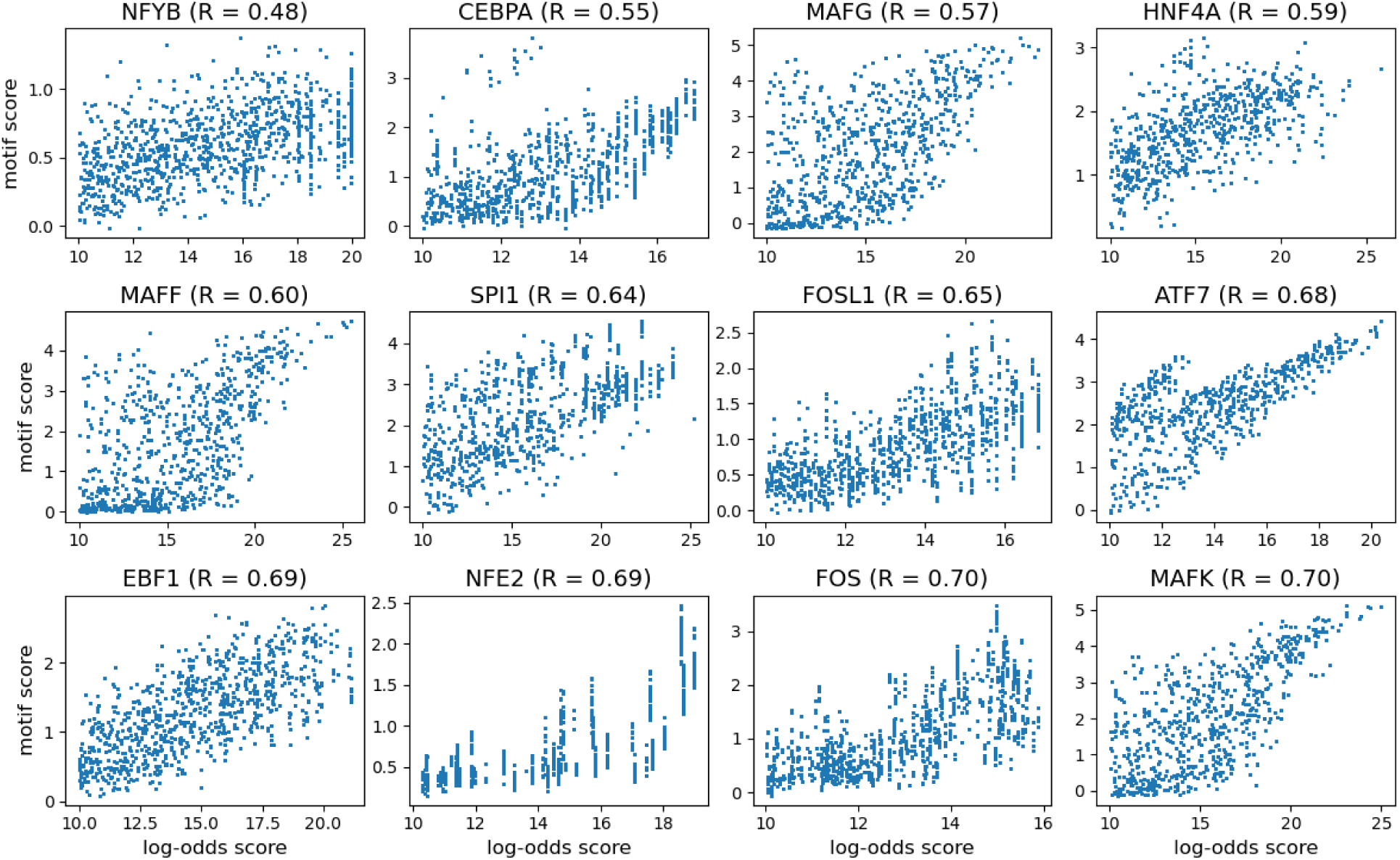
Log-odds score versus Enformer-based motif score for human samples. R indicates Pearson correlation. The 16 samples with lowest correlation are shown.

## References

Arnold, P., Schöler, A., Pachkov, M., Balwierz, P. J., Jørgensen, H., Stadler, M. B., van Nimwegen, E., & Schübeler, D. (2013). Modeling of epigenome dynamics identifies transcription factors that mediate polycomb targeting. Genome research, 23 (1), 60–73.

Arora, S., Yang, J., Akiyama, T., James, D. Q., Morrissey, A., Blanda, T. R., Badjatia, N., Lai, W. K., Ko, M. S., Pugh, B. F., & Mahony, S. (2023). Joint sequence & chromatin neural networks characterize the differential abilities of forkhead transcription factors to engage inaccessible chromatin. *bioRxiv*.

Arvey, A., Agius, P., Noble, W. S., & Leslie, C. (2012). Sequence and chromatin determinants of cell-type–specific transcription factor binding. Genome research, 22 (9), 1723–1734.

Atlasi, Y., Megchelenbrink, W., Peng, T., Habibi, E., Joshi, O., Wang, S.-Y., Wang, C., Logie, C., Poser, I., Marks, H., et al. (2019). Epigenetic modulation of a hardwired 3D chromatin landscape in two naive states of pluripotency. Nature cell biology, 21 (5), 568–578.

Avsec, Ž., Agarwal, V., Visentin, D., Ledsam, J. R., Grabska-Barwinska, A., Taylor, K. R., Assael, Y., Jumper, J., Kohli, P., & Kelley, D. R. (2021). Effective gene expression prediction from sequence by integrating long-range interactions. Nature methods, 18 (10), 1196–1203.

Benjamini, Y., & Speed, T. P. (2012). Summarizing and correcting the GC content bias in high-throughput sequencing. Nucleic acids research, 40 (10), e72–e72.

Berg, O. G., & von Hippel, P. H. (1988). Selection of DNA binding sites by regulatory proteins. Trends in biochemical sciences, 13 (6), 207–211.

Blassberg, R., Patel, H., Watson, T., Gouti, M., Metzis, V., Delás, M. J., & Briscoe, J. (2022). Sox2 levels regulate the chromatin occupancy of WNT mediators in epiblast progenitors responsible for vertebrate body formation. Nature Cell Biology, 24 (5), 633–644.

Boller, S., Ramamoorthy, S., Akbas, D., Nechanitzky, R., Burger, L., Murr, R., Schübeler, D., & Grosschedl, R. (2016). Pioneering activity of the C-terminal domain of EBF1 shapes the chromatin landscape for B cell programming. Immunity, 44 (3), 527–541.

Bulyk, M. L., Drouin, J., Harrison, M. M., Taipale, J., & Zaret, K. S. (2023). Pioneer factors—key regulators of chromatin and gene expression. Nature Reviews Genetics, 24 (12), 809–815.

Chen, X., Xu, H., Yuan, P., Fang, F., Huss, M., Vega, V. B., Wong, E., Orlov, Y. L., Zhang, W., Jiang, J., et al. (2008). Integration of external signaling pathways with the core transcriptional network in embryonic stem cells. Cell, 133 (6), 1106–1117.

Chronis, C., Fiziev, P., Papp, B., Butz, S., Bonora, G., Sabri, S., Ernst, J., & Plath, K. (2017). Cooperative binding of transcription factors orchestrates reprogramming. Cell, 168 (3), 442–459.

Cirillo, L. A., Lin, F. R., Cuesta, I., Friedman, D., Jarnik, M., & Zaret, K. S. (2002). Opening of compacted chromatin by early developmental transcription factors HNF3 (FoxA) and GATA-4. Molecular cell, 9 (2), 279–289.

Dolfini, D., Gatta, R., & Mantovani, R. (2012). NF-Y and the transcriptional activation of CCAAT promoters. Critical reviews in biochemistry and molecular biology, 47 (1), 29–49.

Domcke, S., Bardet, A. F., Adrian Ginno, P., Hartl, D., Burger, L., & Schübeler, D. (2015) Competition between DNA methylation and transcription factors determines binding of NRF1. Nature, 528 (7583), 575–579.

Donaghey, J., Thakurela, S., Charlton, J., Chen, J. S., Smith, Z. D., Gu, H., Pop, R., Clement, K., Stamenova, E. K., Karnik, R., et al. (2018). Genetic determinants and epigenetic effects of pioneer-factor occupancy. Nature genetics, 50 (2), 250–258.

Durdu, S., Iskar, M., Isbel, L., Hoerner, L., Wirbelauer, C., Burger, L., Hess, D., Iesmantavicius, V., & Schübeler, D. (2024). Chromatin-dependent motif syntax defines differentiation trajectories. bioRxiv, 2024–08.

Fidalgo, M., Huang, X., Guallar, D., Sanchez-Priego, C., Valdes, V. J., Saunders, A., Ding, J., Wu, W.-S., Clavel, C., & Wang, J. (2016). Zfp281 coordinates opposing functions of Tet1 and Tet2 in pluripotent states. Cell stem cell, 19 (3), 355–369.

Fu, Y., Sinha, M., Peterson, C. L., & Weng, Z. (2008). The insulator binding protein CTCF positions 20 nucleosomes around its binding sites across the human genome. PLoS genetics, 4 (7), e1000138.

Ghandi, M., Lee, D., Mohammad-Noori, M., & Beer, M. A. (2014). Enhanced regulatory sequence prediction using gapped k-mer features. PLoS computational biology, 10 (7), e1003711.

Gibson, T. J., Larson, E. D., & Harrison, M. M. (2024). Protein-intrinsic properties and context-dependent effects regulate pioneer factor binding and function. Nature Structural & Molecular Biology, 31 (3), 548–558.

Ginno, P. A., Gaidatzis, D., Feldmann, A., Hoerner, L., Imanci, D., Burger, L., Zilbermann, F., Peters, A. H., Edenhofer, F., Smallwood, S. A., et al. (2020). A genome-scale map of DNA methylation turnover identifies site-specific dependencies of DNMT and TET activity. Nature communications, 11 (1), 2680.

Grand, R. S., Burger, L., Gräwe, C., Michael, A. K., Isbel, L., Hess, D., Hoerner, L., Iesmantavicius, V., Durdu, S., Pregnolato, M., et al. (2021). BANP opens chromatin and activates CpG-island-regulated genes. Nature, 596 (7870), 133–137.

Grand, R. S., Pregnolato, M., Baumgartner, L., Hoerner, L., Burger, L., & Schübeler, D. (2024). Genome access is transcription factor-specific and defined by nucleosome position. Molecular Cell.

Hansen, J. L., Loell, K. J., & Cohen, B. A. (2022). A test of the pioneer factor hypothesis using ectopic liver gene activation. Elife, 11, e73358.

Hesselberth, J. R., Chen, X., Zhang, Z., Sabo, P. J., Sandstrom, R., Reynolds, A. P., Thurman, R. E., Neph, S., Kuehn, M. S., Noble, W. S., et al. (2009). Global mapping of protein-DNA interactions in vivo by digital genomic footprinting. Nature methods, 6 (4), 283–289.

Hunter, J. D. (2007). Matplotlib: A 2d graphics environment. Computing in Science & Engineering, 9 (3), 90–95. 10.1109/MCSE.2007.55

Isbel, L., Grand, R. S., & Schübeler, D. (2022). Generating specificity in genome regulation through transcription factor sensitivity to chromatin. Nature Reviews Genetics, 23 (12), 728–740.

Jolma, A., Yin, Y., Nitta, K. R., Dave, K., Popov, A., Taipale, M., Enge, M., Kivioja, T., Morgunova, E., & Taipale, J. (2015). DNA-dependent formation of transcription factor pairs alters their binding specificity. Nature, 527 (7578), 384–388.

Katsuoka, F., & Yamamoto, M. (2016). Small Maf proteins (MafF, MafG, MafK): History, structure and function. Gene, 586 (2), 197–205.

Kelley, D. R. (2020). Cross-species regulatory sequence activity prediction. PLoS computational biology, 16 (7), e1008050.

Khan, A., Fornes, O., Stigliani, A., Gheorghe, M., Castro-Mondragon, J. A., Van Der Lee, R., Bessy, A., Cheneby, J., Kulkarni, S. R., Tan, G., et al. (2018). JASPAR 2018: Update of the open-access database of transcription factor binding profiles and its web framework. Nucleic acids research, 46 (D1), D260–D266.

Kim, S., & Wysocka, J. (2023). Deciphering the multi-scale, quantitative cis-regulatory code. Molecular cell, 83 (3), 373–392.

Korhonen, J., Martinmäki, P., Pizzi, C., Rastas, P., & Ukkonen, E. (2009). MOODS: Fast search for position weight matrix matches in DNA sequences. Bioinformatics, 25 (23), 3181–3182.

Li, M., He, Y., Dubois, W., Wu, X., Shi, J., & Huang, J. (2012). Distinct regulatory mechanisms and functions for p53-activated and p53-repressed DNA damage response genes in embryonic stem cells. Molecular cell, 46 (1), 30–42.

Li, R., Cauchy, P., Ramamoorthy, S., Boller, S., Chavez, L., & Grosschedl, R. (2018). Dynamic EBF1 occupancy directs sequential epigenetic and transcriptional events in B-cell programming. Genes & development, 32 (2), 96–111.

Luo, Z., Gao, X., Lin, C., Smith, E. R., Marshall, S. A., Swanson, S. K., Florens, L., Washburn, M. P., & Shilatifard, A. (2015). Zic2 is an enhancer-binding factor required for embryonic stem cell specification. Molecular cell, 57 (4), 685–694.

Ma, X., Ezer, D., Navarro, C., & Adryan, B. (2015). Reliable scaling of position weight matrices for binding strength comparisons between transcription factors. BMC bioinformatics, 16, 1–13.

Marson, A., Levine, S. S., Cole, M. F., Frampton, G. M., Brambrink, T., Johnstone, S., Guenther, M. G., Johnston, W. K., Wernig, M., Newman, J., et al. (2008). Connecting microRNA genes to the core transcriptional regulatory circuitry of embryonic stem cells. cell, 134 (3), 521–533.

Martin, M. (2011). Cutadapt removes adapter sequences from high-throughput sequencing reads. *EMBnet*. journal, 17 (1), 10–12.

Mayran, A., Khetchoumian, K., Hariri, F., Pastinen, T., Gauthier, Y., Balsalobre, A., & Drouin, J. (2018). Pioneer factor Pax7 deploys a stable enhancer repertoire for specification of cell fate. Nature Genetics, 50 (2), 259–269.

Michael, A. K., Grand, R. S., Isbel, L., Cavadini, S., Kozicka, Z., Kempf, G., Bunker, R. D., Schenk, A. D., Graff-Meyer, A., Pathare, G. R., et al. (2020). Mechanisms of OCT4-SOX2 motif readout on nucleosomes. Science, 368 (6498), 1460–1465.

Minderjahn, J., Schmidt, A., Fuchs, A., Schill, R., Raithel, J., Babina, M., Schmidl, C., Gebhard, C., Schmidhofer, S., Mendes, K., et al. (2020). Mechanisms governing the pioneering and redistribution capabilities of the non-classical pioneer PU.1. Nature communications, 11 (1), 402.

Mirny, L. A. (2010). Nucleosome-mediated cooperativity between transcription factors. Proceedings of the National Academy of Sciences, 107 (52), 22534–22539.

Oldfield, A. J., Yang, P., Conway, A. E., Cinghu, S., Freudenberg, J. M., Yellaboina, S., & Jothi, R. (2014). Histone-fold domain protein NF-Y promotes chromatin accessibility for cell type-specific master transcription factors. Molecular cell, 55 (5), 708–722.

Ostapcuk, V., Mohn, F., Carl, S. H., Basters, A., Hess, D., Iesmantavicius, V., Lampersberger, L., Flemr, M., Pandey, A., Thomä, N. H., et al. (2018). Activity-dependent neuroprotective protein recruits HP1 and CHD4 to control lineage-specifying genes. Nature, 557 (7707), 739–743.

Pagès, H., Aboyoun, P., Gentleman, R., & DebRoy, S. (2024). Biostrings: Efficient manipulation of biological strings [R package version 2.74.1]. 10.18129/B9.bioc.Biostrings

Pedregosa, F., Varoquaux, G., Gramfort, A., Michel, V., Thirion, B., Grisel, O., Blondel, M., Prettenhofer, P., Weiss, R., Dubourg, V., et al. (2011). Scikit-learn: Machine learning in Python. the Journal of machine Learning research, 12, 2825–2830.

Peng, Y., Song, W., Teif, V. B., Ovcharenko, I., Landsman, D., & Panchenko, A. R. (2024). Detection of new pioneer transcription factors as cell-type-specific nucleosome binders. Elife, 12, RP88936.

Pham, T.-H., Minderjahn, J., Schmidl, C., Hoffmeister, H., Schmidhofer, S., Chen, W., Längst, G., Benner, C., & Rehli, M. (2013). Mechanisms of in vivo binding site selection of the hematopoietic master transcription factor PU.1. Nucleic acids research, 41 (13), 6391–6402.

Pique-Regi, R., Degner, J. F., Pai, A. A., Gaffney, D. J., Gilad, Y., & Pritchard, J. K. (2011). Accurate inference of transcription factor binding from DNA sequence and chromatin accessibility data. Genome research, 21 (3), 447–455.

Pop, R. T., Pisante, A., Nagy, D., Martin, P. C., Mikheeva, L. A., Hayat, A., Ficz, G., & Zabet, N. R. (2023). Identification of mammalian transcription factors that bind to inaccessible chromatin. Nucleic Acids Research, 51 (16), 8480–8495.

Scelfo, A., Fernández-Pérez, D., Tamburri, S., Zanotti, M., Lavarone, E., Soldi, M., Bonaldi, T., Ferrari, K. J., & Pasini, D. (2019). Functional landscape of PCGF proteins reveals both RING1A/B-dependent-and RING1A/B-independent-specific activities. Molecular cell, 74 (5), 1037–1052.

Seabold, S., & Perktold, J. (2010). Statsmodels: Econometric and statistical modeling with python. 9th Python in Science Conference.

Sherwood, R. I., Hashimoto, T., O’donnell, C. W., Lewis, S., Barkal, A. A., Van Hoff, J. P., Karun, V., Jaakkola, T., & Gifford, D. K. (2014). Discovery of directional and nondirectional pioneer transcription factors by modeling DNase profile magnitude and shape. Nature biotechnology, 32 (2), 171–178.

Shrikumar, A., Greenside, P., & Kundaje, A. (2017). Learning important features through propagating activation differences. International conference on machine learning, 3145–3153.

Soufi, A., Donahue, G., & Zaret, K. S. (2012). Facilitators and impediments of the pluripotency reprogramming factors’ initial engagement with the genome. Cell, 151 (5), 994–1004.

Soufi, A., Garcia, M. F., Jaroszewicz, A., Osman, N., Pellegrini, M., & Zaret, K. S. (2015). Pioneer transcription factors target partial DNA motifs on nucleosomes to initiate reprogramming. Cell, 161 (3), 555–568.

Srivastava, D., Aydin, B., Mazzoni, E. O., & Mahony, S. (2021). An interpretable bimodal neural network characterizes the sequence and preexisting chromatin predictors of induced transcription factor binding. Genome biology, 22, 1–25.

Stadler, M. B., Murr, R., Burger, L., Ivanek, R., Lienert, F., Schöler, A., Nimwegen, E. v., Wirbelauer, C., Oakeley, E. J., Gaidatzis, D., et al. (2011). DNA-binding factors shape the mouse methylome at distal regulatory regions. Nature, 480 (7378), 490–495.

Stoeber, S., Godin, H., Xu, C., & Bai, L. (2024). Pioneer factors: Nature or nurture Critical Reviews in Biochemistry and Molecular Biology, 1–15.

Tan, G. (2017). JASPAR2018: Data package for JASPAR 2018 [R package version 1.1.1]. http://jaspar.genereg.net/

Tareen, A., & Kinney, J. B. (2020). Logomaker: Beautiful sequence logos in python. Bioinformatics, 36 (7), 2272–2274.

Teng, M., & Irizarry, R. A. (2016). Accounting for GC-content bias reduces systematic errors and batch effects in ChIP-seq peak callers. bioRxiv, 090704.

Teytelman, L., Thurtle, D. M., Rine, J., & van Oudenaarden, A. (2013). Highly expressed loci are vulnerable to misleading ChIP localization of multiple unrelated proteins. Proceedings of the National Academy of Sciences, 110 (46), 18602–18607.

Thanos, D., & Maniatis, T. (1995). Virus induction of human IFN*β* gene expression requires the assembly of an enhanceosome. Cell, 83 (7), 1091–1100.

Wang, J., Zhuang, J., Iyer, S., Lin, X., Whitfield, T. W., Greven, M. C., Pierce, B. G., Dong, X., Kundaje, A., Cheng, Y., et al. (2012). Sequence features and chromatin structure around the genomic regions bound by 119 human transcription factors. Genome research, 22 (9), 1798–1812.

Waskom, M. L. (2021). Seaborn: Statistical data visualization. Journal of Open Source Software, 6 (60), 3021. 10.21105/joss.03021

Xu, C., Kleinschmidt, H., Yang, J., Leith, E. M., Johnson, J., Tan, S., Mahony, S., & Bai, L. (2024). Systematic dissection of sequence features affecting binding specificity of a pioneer factor reveals binding synergy between FOXA1 and AP-1. Molecular cell, 84 (15), 2838–2855.

Yan, J., Enge, M., Whitington, T., Dave, K., Liu, J., Sur, I., Schmierer, B., Jolma, A., Kivioja, T., Taipale, M., et al. (2013). Transcription factor binding in human cells occurs in dense clusters formed around cohesin anchor sites. Cell, 154 (4), 801–813.

Yin, Y., Morgunova, E., Jolma, A., Kaasinen, E., Sahu, B., Khund-Sayeed, S., Das, P. K., Kivioja, T., Dave, K., Zhong, F., et al. (2017). Impact of cytosine methylation on DNA binding specificities of human transcription factors. Science, 356 (6337), eaaj2239.

Yue, F., Cheng, Y., Breschi, A., Vierstra, J., Wu, W., Ryba, T., Sandstrom, R., Ma, Z., Davis, C., Pope, B. D., et al. (2014). A comparative encyclopedia of DNA elements in the mouse genome. Nature, 515 (7527), 355–364.

Zhang, Y., Liu, T., Meyer, C. A., Eeckhoute, J., Johnson, D. S., Bernstein, B. E., Nusbaum, C., Myers, R. M., Brown, M., Li, W., et al. (2008). Model-based analysis of ChIP-seq (MACS). Genome biology, 9, 1–9.

